# Frequent assembly of chimeric complexes in the protein interaction network of an interspecies yeast hybrid

**DOI:** 10.1101/2020.06.02.130567

**Authors:** Rohan Dandage, Caroline M. Berger, Isabelle Gagnon-Arsenault, Kyung-Mee Moon, Richard Greg Stacey, Leonard J. Foster, Christian R. Landry

**Affiliations:** Département de Biochimie, Microbiologie et Bio-informatique, Faculté des sciences et de génie, Université Laval, Québec, Québec, G1V 0A6, Canada; PROTEO, Le réseau québécois de recherche sur la fonction, la structure et l’ingénierie des protéines, Université Laval, Québec, Québec, G1V 0A6, Canada; Centre de Recherche en Données Massives (CRDM), Université Laval, Québec, Québec, G1V 0A6, Canada; Département de Biologie, Faculté des sciences et de génie, Université Laval, Québec, Québec, G1V 0A6, Canada; Department of Biochemistry & Molecular Biology, and Michael Smith Laboratories, University of British Columbia, Vancouver, British Columbia, V6T 1Z4, Canada.

## Abstract

Hybrids between species often show extreme phenotypes, including some that take place at the molecular level. In this study, we investigated the phenotypes of an interspecies diploid hybrid in terms of protein-protein interactions inferred from protein correlation profiling. We used two yeast species, *Saccharomyces cerevisiae* and *Saccharomyces uvarum*, which are interfertile, but yet have proteins diverged enough to be differentiated using mass spectrometry. Most of the protein-protein interactions are similar between hybrid and parents, and are consistent with the assembly of chimeric complexes, which we validated using an orthogonal approach for prefoldin complex. We also identify instances of altered protein-protein interactions in the hybrid, for instance in complexes related to proteostasis and in mitochondrial protein complexes. Overall, this study uncovers likely frequent occurrence of chimeric protein complexes with few exceptions, which may result from incompatibilities or imbalances between the parental proteins.

## Introduction

One of the major goals of evolutionary biology is to understand the molecular basis of phenotypic diversity, which fuels evolution by natural selection. Hybrids between species provide a unique opportunity to investigate the molecular underpinnings of phenotypic diversity. One could expect hybrids to show an intermediate phenotype, in which their characters are mid-way between parental phenotypes. However, hybrids often show a large spectrum of phenotypes, including extreme values that are unexpected given the parental traits (Rieseberg et al. 1999; Landry et al. 2007; Maheshwari & Barbash 2011; Bar-Zvi et al. 2017). Indeed, previous studies have demonstrated the importance of large-scale regulatory rewiring in hybrids that impacts many processes and molecular phenotypes such as nucleosome positioning, translation efficiency, protein abundance, methylation, transcription of non-coding RNA, and the replication program (Landry et al. 2007; Tirosh et al. 2010; Tirosh & Barkai 2011; McManus et al. 2014; Zhu et al. 2017; Bar-Zvi et al. 2017; Bamberger et al. 2018; Zhao et al. 2018).

In the context of a cell, protein-protein interactions (PPIs) are central to molecular functions. Stable, non-transient PPIs lead to the formation of protein complexes with diverse functions such as DNA replication, repair and transcription, transport, catalysis, signaling and many others (Sowmya et al. 2015). Since protein complexes have such an important role, variation in their organization impacts the phenotype of organisms. Therefore, examining how PPI networks integrate in interspecific hybrids is of great interest.

Variation in the composition of protein complexes caused by hybridization between species can lead to many qualitative outcomes (as shown in Figure 1A). Since hybrids between species that are amenable to genetic studies often do not suffer from dramatic fitness loss, we can broadly expect proteins to interact in the hybrids as they do in the parental species for complexes related to core cellular functions. The complexes could then be of two types: chimeric and parental. Chimeric complexes would result from interlogous (interspecies) and intralogous (intraspecies) PPIs and therefore, they would be a mixture of the two parental proteomes. On the other hand, some parental complexes could assemble from proteins of one parental species only, i.e. complexes would be formed only or majoritarily by intralogous PPIs. Parental complexes in hybrid could preferentially form, for instance, if affinities are higher among proteins from the same parental species than between species, or if difference in timing of expression in cis between species prevents interlogous PPIs. Lastly, another possibility is that novel complexes could emerge in the hybrids, i.e. complexes with interlogous PPIs that do not have equivalent PPIs in the parents.

**Figure 1.**
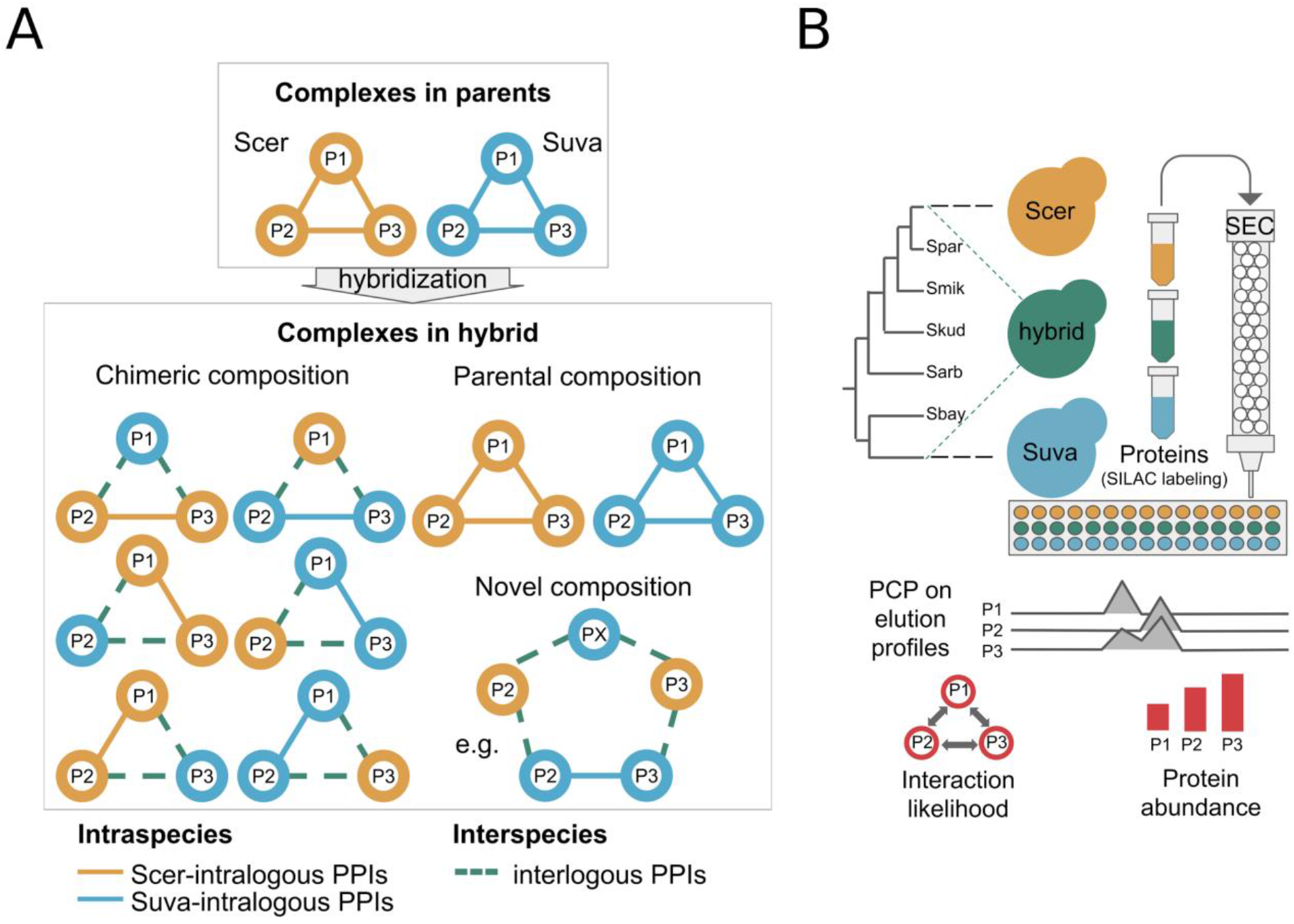
Possible scenarios for the assembly of a protein complex in hybrid and experimental layout of the comparative proteomics approach. **A)** Scenarios depicting the assembly of a three-protein complex (P1, P2 and P3) in hybrid, with intralogous and interlogous PPIs. **B)** Experimental layout of the comparative proteomics approach. In the first step, proteins of hybrid and parental strains were labeled with different lysine isotopes (see Methods and Figure S2 for details). Here, parental strains were used as controls. After extraction, proteins were separated by Size-Exclusion Chromatography (SEC) and the eluted fractions were processed by mass spectrometry. Elution profiles of the proteins were used to infer PPIs and protein complexes by Protein Correlation Profiling (PCP), using Dynamic Time Warping (DTW) method (Figure S3), with the principle that two proteins that interact would tend to have similar elution profiles. The peptide counts per protein were used to estimate protein abundance in hybrid and parental species (See methods). In the figure, we schematically represent the elution profiles of hypothetical proteins P1, P2 and P3. Using the correlations between their elution profiles and peptide counts per protein, the likelihood of PPIs and protein abundances were respectively estimated.

The null hypothesis that protein complexes appear to assemble in hybrids as they do in parents is supported by previous studies. For instance, the assembly of the nuclear pore complex and of the RNA polymerase II in yeast hybrids are consistent with the conservation of complexes in hybrids (Leducq et al. 2012), i.e. interlogous PPIs are formed in hybrids in a way that reflects the parental PPIs. In fact, it is possible that most proteins between closely related species are capable of interacting this way given that proteins that are as diverged as those of yeast and humans can complement and interact with each other (Kachroo et al. 2015; Zhong et al. 2016). However, cases of incompatibilities and of novel interactions do exist. For instance, a study supported the scenario whereby proteins from two species form only intralogous PPIs when expressed in the same cell and co-evolution within species of the parental proteins belonging to the PCNA (Proliferating Cell Nuclear Antigen) complex prevents interlogous PPIs (Zamir et al. 2012). On the other hand, another study found that some complexes contain interlogous PPIs that are not seen in parental species. They further linked one of these new hybrid complexes to an enhanced function in tryptophan transport (Piatkowska et al. 2013). Both deleterious and adaptive changes can thus occur in hybrid complexes.

Because previous studies were limited in terms of coverage of molecular functions, whether hybridization is associated with a global reorganization of protein complexes in the cell is still mostly unexplored. Until recently, we lacked experimental methods that allow studying of a large number of protein complexes simultaneously in hybrids and parental species. Here, we applied a method that allows us to survey several protein complexes simultaneously to a yeast hybrid. Several reasons make yeast species excellent experimental models to address the question as to how protein complexes assemble in hybrids. Firstly, hybrids can be readily produced in the laboratory (Krogerus et al. 2018). Second, spontaneous hybridization is common among yeast species and may have an important impact on their performance in nature (Leducq et al. 2016; Barbosa et al. 2016). Third, during industrial processes such as fermentation, hybridization is thought to be an important mechanism for adaptation (González et al. 2006; López-Malo et al. 2013). For example, hybridization leads to a high fermentation capacity with a desirable aroma profile for beer fermentations (Mertens et al. 2015). And finally, the yeast proteome and PPI network have been well characterized (Ito et al. 2001; Krogan et al. 2006; Tarassov et al. 2008), making it an ideal model for species comparisons.

We applied Size-Exclusion Chromatography (SEC), Protein Correlation Profiling (PCP) and Stable Isotope Labeling by Amino acids in Cell culture (SILAC), SEC-PCP-SILAC (Kristensen & Foster 2014), to the budding yeast *Saccharomyces cerevisiae* (denoted as Scer), to *Saccharomyces uvarum* (denoted as Suva) (Scannell et al. 2011) and to their F1 diploid hybrid. SEC-PCP-SILAC separates complex mixtures of endogenous proteins into a set of fractions that are analysed by mass spectrometry. The mass spectrometry method measured the abundance of ~400 proteins, with a reasonably broad coverage of molecular functions of the proteomes. The profiles of co-migrating proteins were clustered to reconstruct PPIs and thus protein complexes (Kristensen et al. 2012). From this method, we inferred likelihoods of PPIs, which we used in the comparative analysis between hybrid and parental species.

## Results and discussion

### Proteomics approach to monitor PPIs in hybrid and parental strains

Scer and Suva are among the most divergent species in the *Saccharomyces* phylogeny (average distance of 50 million years (Kellis et al. 2003)), with an overall nucleotide sequence divergence between 30-35% (Morales & Dujon 2012) and a 16% divergence at the protein level (Figure S1A). Scer and Suva strains were used as parental species along with their diploid F1 progeny as hybrid produced in two biological replicates (Figure 1B and Table S1A for genotypes of the strains). The proteomes of the two parental strains are divergent enough that peptides could in general be distinguished by mass spectrometry (median Jaccard distance of 82.1%, Figure S1B). The proteomics approach of SEC-PCP-SILAC allowed us to resolve likely interacting proteins (using SEC) and to infer putative PPIs (using PCP), in a comparative manner (using SILAC) (Figure 1B, Figure S2 and see Methods for details).

Because some peptides derived from conserved sequences of orthologous proteins cannot be assigned to one parental species or the other in hybrid, only uniquely aligned peptides that could be ascribed to a protein of either Scer or Suva were used in the analysis. The uniquely aligned peptides amount to ~70% of the total number of peptides detected (Figure S1C), representing ~80% (~400) of the total number of proteins detected (Figure S1D). Since some proteins were detected by uniquely as well as non-uniquely aligned peptides, only a relatively small fraction of the total number of proteins were removed by the filtering of proteins that were detected exclusively by non-uniquely aligned peptides. Thus, there was not an undue bias towards high sequence divergence in the remaining set of proteins (Figure S1A, Data S1). This allowed a larger coverage of molecular functions, and not only for the proteins that diverge the most between species. Finally, in the PCP step, the elution spectra of proteins were correlated with each other by Dynamic Time Warping (DTW, see Methods). Similar elution profiles indicate likely interaction between a given pair of proteins. Therefore, pairwise similarity estimates between all protein pairs of a species provided a putative PPI network for that species (Data S2).

The use of uniquely aligned peptides allowed us to differentiate intralogous and interlogous PPIs with high confidence. In contrast to intralogous PPIs which can occur in parent as well as in the hybrid, interlogous PPIs can only take place in the hybrid. Therefore, for the comparative analysis, as a reference for interlogous PPIs, we inferred the expected interlogous PPIs from the elution profiles obtained for the parental species. Specifically, the elution profiles from the parental species were treated as if they originated from the same cell, called a ‘theoretical hybrid’. Then, such elution profiles were correlated by DTW to obtain putative PPIs which we hereon refer to as ‘expected interlogous PPIs’.

Since for the parental Scer, protein abundance and PPIs are well characterized, we used them as references to assess the accuracy of the SEC-PCP-SILAC data. In order to estimate the likelihoods of PPIs between pairs of proteins, we applied DTW to calculate the distance between elution profiles of the proteins. Using the elution profiles of Scer proteins, we first scanned through a range of window sizes used for DTW and a range of thresholds applied on the distance between pairs of elution profiles to obtain optimal parameters (Figure S3, see Methods) that provide a best match to the known PPIs from databases (Figure 2A). Also, upon comparing the protein abundance scores obtained from our proteomics data (Data S3) with the reference proteomics data obtained from PAXdb (Wang et al. 2015), we obtained a significant positive correlation (Figure 2B). Overall, for both the likelihood of PPIs and protein abundance, the Scer proteomics data show high similarity with known reference values for Scer in databases.

**Figure 2.**
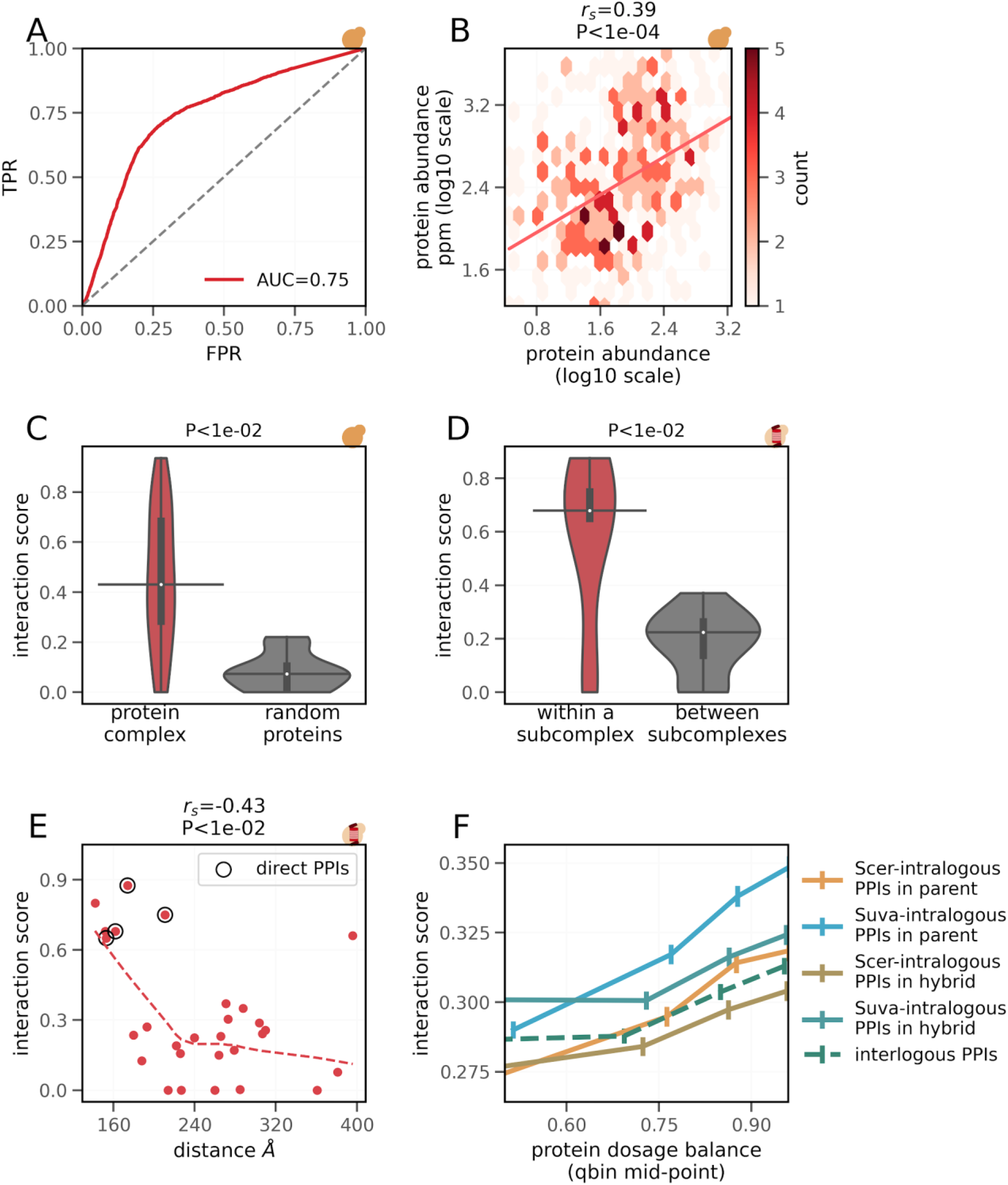
Primary validation of the proteomics data. SEC-PCP-SILAC data for parental Scer species was compared against reference data obtained from databases (panels A to E). The panels showing the analysis with parental Scer strain (**A, B** and **C**) are annotated with the logo of Scer, while those showing the analysis with proteasome complex in parental Scer strain (panel **D** and **E**) are annotated with the logo representing a proteasome inside Scer. **A)** Comparison of experimentally determined interaction scores with the association of interactors in protein complexes obtained from databases (see Methods). AUC: Area Under Curve, TPR: True Positive Rate and FPR: False Positive Rate. **B)** Comparison of experimentally determined protein abundance (shown on x-axis) with reference protein abundance data obtained from proteomics source of PeptideAtlas (Desiere et al. 2006) from PAXdb (Wang et al. 2015). r_s_: Spearman’s correlation coefficient. The colorbar indicates the count of proteins in the hexbins. **C)** Comparison of interaction scores among members of protein complexes and among random sets of proteins (of equal size and number as for protein complexes) that are not part of any protein complex from parental Scer species. P-value from two-sided Mann-Whitney U tests is shown. On the violin plots, the medians of the distributions are shown by a horizontal black line and quartiles by a vertical thick black line. **D)** Comparison of interaction scores within and between subcomplexes of the proteasome. Subcomplex-wise lists of proteins were obtained from Saccharomyces Genome Database (SGD) (Cherry et al. 2012). P-value from two-sided Mann-Whitney U tests is shown. On the violin plots, the medians of the distributions are shown by a horizontal black line and quartiles by a vertical thick black line. **E)** Correlation between the interaction scores of the proteasome complex of Scer and their inter-protein distances. Dashed line shows the fitted values from a lowess regression. Known direct PPIs within the proteasome complex were obtained from the BioGRID database (Chatr-Aryamontri et al. 2017). Inter-protein distances in the complex were obtained from Chrétien et al. (Chrétien et al. 2018). r_s_: Spearman’s correlation coefficient. **F)** Relationship between dosage balance and interaction score. For clarity, dosage balance values are divided into bins of equal size (on x-axis). Average interaction score per bin is shown (on y-axis) with bars indicating 95% confidence interval. Dosage balance is measured as `1-((|p1-p2|)/(p1+p2)),` where `p1` and `p2` are protein abundances of two proteins.

In order to further assess if the inferred PPIs are indicative of the association between proteins, we compared the average interaction scores for known protein complexes (obtained from Complex Portal database (Meldal et al. 2019)) from Scer data with randomly drawn sets of proteins that are not known to be part of any protein complexes (of equal size and number as for protein complexes) (Figure 2C). The interaction scores for protein complexes are significantly higher than the ones for random sets of proteins (P<1e-2, two-sided Mann-Whitney U test), indicating that interaction scores indeed capture associations among proteins. For example, in the Scer proteasome, we find significantly higher interaction scores for PPIs within subcomplexes, compared to PPIs between subcomplexes (P<1e-2, two-sided Mann-Whitney U test, as shown in Figure 2D. See Figure S4 for PPI network within proteasome complex). We also find that interaction scores significantly correlate with the inter-protein distances within the complex (Figure 2E), indicating that empirically obtained interaction scores, i.e. likelihoods of PPIs, are sensitive to the physical distances between the proteins and thus, that they are a good proxy for association between proteins. Moreover, this analysis for the proteasome protein complex also suggests that the higher values of interaction scores are likely to indicate direct PPIs.

Since the abundance of interacting proteins is often balanced within protein complexes (Ge et al. 2001; Taggart & Li 2018), we tested, for all comparisons, if the likelihoods of PPIs are positively related with the dosage balance of proteins. From the proteomics data, we captured an expected positive relationship for almost all intralogous PPIs(Figure 2F). Following the relationship between the interaction score and dosage balance, we found that interactors within complexes (annotated for Scer) show significantly higher interaction scores than interactors that are not contained in a protein complex (Figure S5A). They also maintain greater dosage balance within complexes (Figure S5B), corroborating with earlier reports (Ge et al. 2001; Veitia et al. 2008; Ishikawa et al. 2017). Overall, the biological significance of the empirical data, as revealed from its conformity with benchmarks and other known trends, lead us to the further comparative analysis of PPIs between hybrid and parental species.

### Strong similarity between parental and hybrid PPIs and for chimeric complexes in hybrid

We compared the scores of intralogous and interlogous PPIs to examine how PPIs form in the hybrid relative to parental species. In each case, we estimated the overall extent of the similarity between PPIs using rank correlation coefficients (Figure 3A). Intralogous PPIs in hybrid show greater similarity with the intralogous PPIs in the parents (Figure 3A, left). This trend holds true for both Scer and Suva intralogous PPIs (Figure 3A, middle) and even in the case of interlogous PPIs relative to the expected interlogous PPIs in the theoretical hybrid (Figure 3A, right). In support to this analysis, a similar trend of conservation was observed upon clustering of interaction scores and protein abundance (Figure S6). Interestingly, intralogous PPIs in the hybrid are more similar to those in the parents than for any other comparisons, suggesting that pairs of proteins from the same species tend to interact similarly in hybrids, as they do in the parents. This may reflect a slight preference for intralogous PPIs in hybrids. However, as discussed below, we find limited additional support for this.

**Figure 3.**
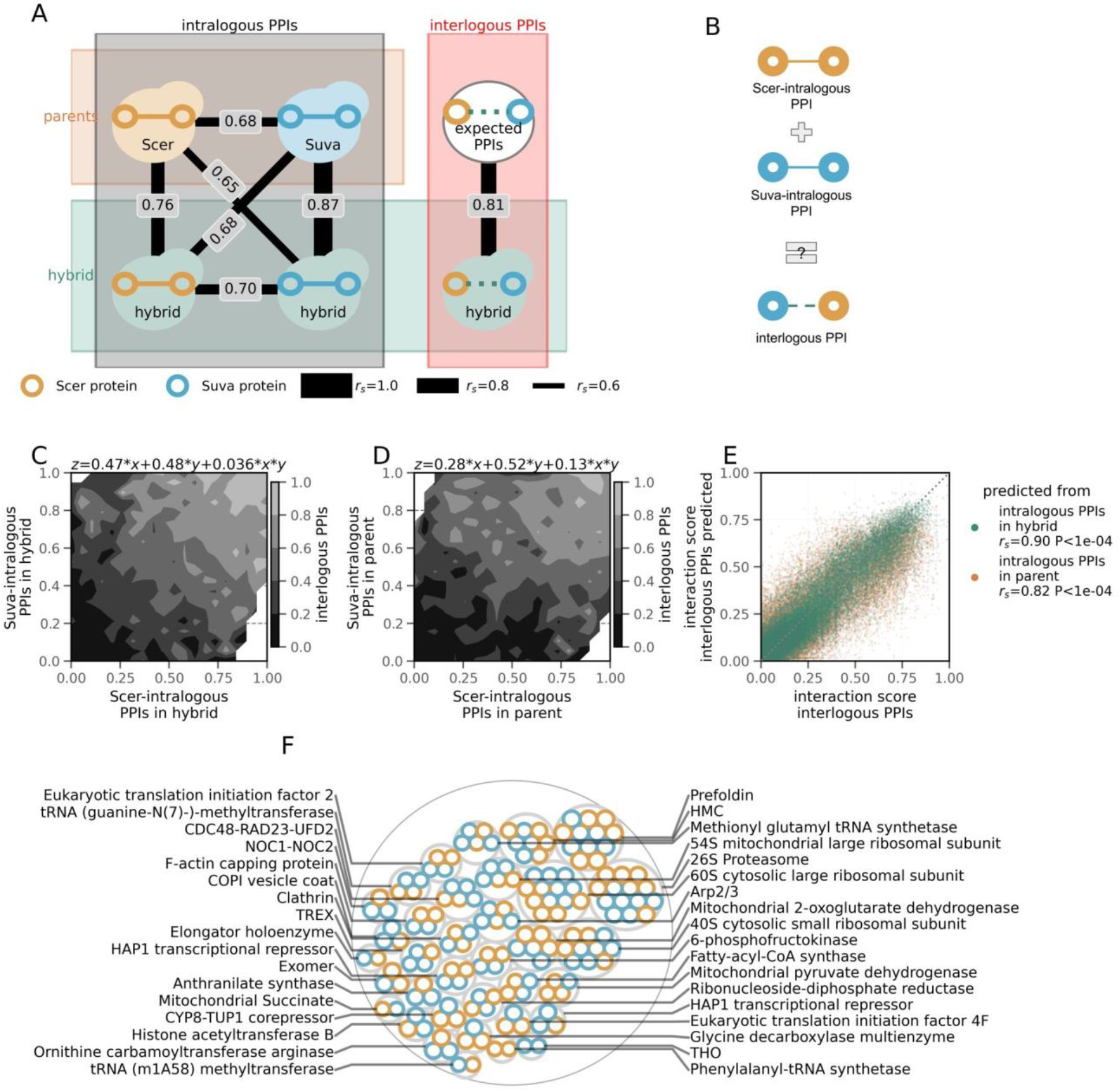
General conservation of PPIs in hybrids and formation of chimeric complexes. **A)** Putative PPIs in hybrids and parents were classified as intralogous if both interactors belong to the same species and as interlogous otherwise. Similarity of interaction scores (estimated by Spearman’s correlation coefficient r(_s_)) between these classes of PPIs in hybrids and parents (see text) is shown. The thickness of the lines connecting the classes is in scale with r_s_. **B)** Schematic representing the model in which likelihood of interlogous PPIs in hybrid is an additive effect of the intralogous PPIs. **C)** Contour plot representing the relationship between the interaction scores in the hybrid of Scer-intralogous (on x-axis) and Suva-intralogous PPIs (on y-axis), with that of interlogous PPIs (on z-axis). The fitting equation of multiple linear regression between the x and y variables is shown above the plot. The interaction scores of two possible interlogous PPIs between two interactors (each from a species) were averaged as noted in the text. **D)** Similar plot as in C, except that interaction scores of intralogous PPIs in the parents instead of the hybrid are lined on x- and y-axes. **E)** The correlations between the empirically estimated interaction scores for interlogous PPIs in hybrid (x-axis) and the interaction scores of intralogous PPIs predicted by multiple linear regression from interaction scores of intralogous PPIs in parents (orange) or in hybrid (green). r_s_: Spearman’s correlation coefficient. **F)** Protein complexes detected in the hybrid. Scer proteins are highlighted with a yellow border, while Suva proteins are highlighted with a blue border.

In order to further quantify the similarity of the hybrid PPIs compared with the parental ones, we tested whether the likelihoods of intralogous PPIs in parents can predict the likelihood of interlogous PPIs in hybrid (Figure 3B). This analysis considers that the likelihood of an interlogous PPI is the average of PPI scores of two possible interlogous PPIs, i.e. interaction score between P1 of Scer with P2 of Suva and interaction score between P2 of Scer with P1 of Suva are averaged. The positive relationship between the likelihoods of interlogous PPIs in the hybrid with likelihoods of intralogous PPIs in the hybrid (Figure 3C) and in parents (Figure 3D) is clearly evident on the contour maps. Moreover, the fitting of multiple linear regression suggests that the likelihoods of intralogous PPIs contribute equally towards predicting the likelihood of interlogous PPIs. We obtain a stronger association for prediction from intralogous PPIs in hybrid compared to that from intralogous PPIs in parents (Figure 3E), which is expected if the PPI profile in hybrids is overall slightly different from the parental ones. These results indicate that the likelihood of interlogous PPIs in hybrid can at least partially be additively predicted from the likelihoods of intralogous PPIs, either in the parents or in the hybrid, again supporting an overall conservation of PPIs in hybrids.

We next looked directly at known protein complexes that have been characterized in Scer. We considered protein complexes for which at least one PPI was captured in the proteomics data, resulting in 35 protein complexes in the hybrid (Figure 3F), 21 in Scer parent and 25 in Suva parent. The variation in the number of protein complexes detected is due to the variation in the number of proteins detected and of PPIs inferred in each case. The smaller number of protein complexes in parental Scer and Suva species is caused by the smaller number of proteins detected (405 Scer proteins and 401 Suva proteins) as compared to the hybrid (418 Scer proteins and 421 Suva proteins) which consequently led to a relatively smaller number of PPIs inferred in parental species (intralogous PPIs only) as compared to the hybrid (intralogous and interlogous PPIs). In the hybrid, the high likelihoods of interlogous PPIs suggest frequent assembly of chimeric complexes (e.g. in proteasome and prefoldin complexes as shown in Figure S7 and S8 respectively). Assessing the relative proportion of complexes with a chimeric (those that contain interlogous PPIs) versus a parental composition (those that contain only intralogous PPIs) is challenging (Figure 1A). Indeed, the number of possible chimeric assemblies of protein complexes in the hybrid increases exponentially with the number of subunits because each position in a complex can be occupied by either one of the two parental proteins. To assess the proportion of complexes in these two categories, we considered likelihoods of interlogous PPIs in hybrid as a proxy for the number of protein complexes with chimeric composition, and likelihoods of intralogous PPIs in hybrid as a proxy for protein complexes with parental composition. Because protein complexes with chimeric composition would also contain intralogous PPIs, the proxy for the number of protein complexes with chimeric composition is an underestimate.

Using median interaction scores as a threshold, we segmented each type of PPI into ‘low’ (interaction score < median) and ‘high’ (interaction score > median) likelihood. Here, the protein complexes in the ‘high’ class are considered to assemble in hybrid. A similar number of complexes are represented in the two categories (Figure S9), suggesting that protein complexes with chimeric composition are generally at least as abundant as those with parental composition in the hybrid. This trend supports the absence of significant bias in the preference for either intralogous or interlogous or PPIs in general (Figure S10). Considering that the proxy for the number of chimeric protein complexes was an underestimate, the chimeric protein complexes are extremely likely to be more frequent in the hybrid than pure parental complexes.

We also assessed the presence of putative novel complexes in hybrid by clustering the PPIs with significantly high interaction scores in hybrid but significantly low interaction scores in parents. This analysis revealed a set of proteins, consisting of 4 Suva proteins (Ahp1, Pcm1, Scw4 and Tsa1) and a Scer protein (Adh4), that do not have any overlap with any known protein complexes. The molecular function of this putative protein complex is challenging to infer. It might be related to thioredoxin peroxidase activity as two of the five Suva proteins (Ahp1 and Tsa1) are involved in that molecular function. Interestingly, however, all the subunits of protein complexes are not known to colocalize with each other. Therefore, this evidence for the possibility of formation of a novel complex needs further experimental validation. Overall, the lack of power in the data limited in depth analysis in the direction of identification of novel protein complexes.

One of the complexes that appears to be chimeric based on these analyses is the prefoldin complex, which we next used as a model to validate the chimeric complexes. (Figure S8). Prefoldin is a hexameric protein complex that acts as a molecular co-chaperone involved in cytoplasmic folding of actin and tubulin monomers during cytoskeleton assembly (Millán-Zambrano and Chávez 2014). We probed PPIs between pairs of proteins in the prefoldin complex using Dihydrofolate Reductase Protein-Fragment Complementation Assay (DHFR-PCA, see Figure 4A for experimental layout, and Figure S11 and Methods for more details) (Tarassov et al. 2008; Michnick et al. 2010; Freschi et al. 2013). DHFR-PCA provides a quantitative signal of interaction strength for direct and near direct physical interactions (Schlecht et al. 2012; Freschi et al. 2013; Levy et al. 2014; Diss et al. 2017; Diss & Lehner 2018), allowing quantitative comparisons with the proteomics data. From the DHFR-PCA experiments, we obtained relative strengths of PPIs (Data S4) within the complex in the hybrid (Figure 4B) and in parents (Figure S12).

**Figure 4.**
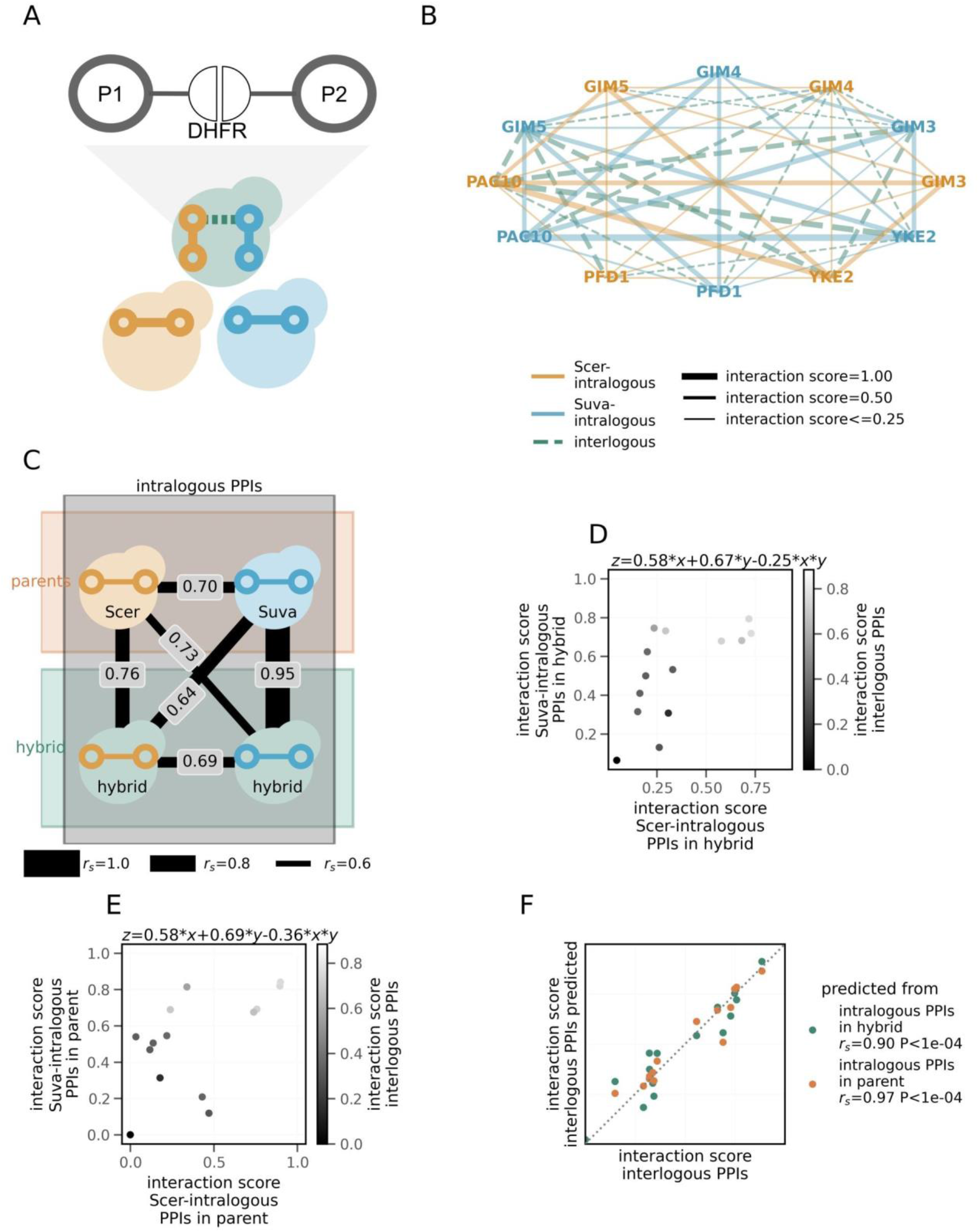
Orthogonal validation for the presence of chimeric PPIs in the prefoldin complex. **A)** Experimental layout of the DHFR-PCA method. Individual yeast strains harboring different combinations of two interacting proteins in the prefoldin complex (each protein tagged with a complementary fragment of the DHFR) (see Tables S1A and S1C, and Methods for details) allowed testing for pairwise PPIs: intralogous in parents and both intralogous and interlogous in hybrid. **B)** PPI network within the prefoldin complex in hybrid, obtained from DHFR-PCA. **C)** Similarity of intralogous and interlogous interactions for interaction strengths in the prefoldin complex detected by DHFR-PCA. **D)** Relationship between the strength of intralogous and interlogous PPIs in the hybrid. Color of the dots is scaled according to the interaction score, i.e. the strength of interlogous PPI in hybrid. The fitted multiple linear regression equation is shown at the top of the plot. **E)** Similar analysis as in panel D, except that the interaction scores of the intralogous PPIs in parental species are lined on the x and y-axis. **F)** Correlation between the actual strength of interlogous PPIs in hybrid with the ones predicted from intralogous PPIs in hybrid and in parents. Spearman’s correlation coefficient (rs) is denoted for each case in the plot legend.

Because the DHFR-PCA experiment detected physical PPIs within complexes between Scer and Suva proteins in the hybrid, it confirms the existence of interlogous complex assembly. Also, the strength of the interlogous PPIs in hybrid (Figure 4B) confirms the chimeric composition of the prefoldin complex observed above. As generally observed in the proteomics data (Figure 3A), intralogous PPI scores in parents are more similar to the intralogous PPIs in the hybrid than when comparing the two parental species to each other (Figure 4C). Furthermore, we tested if likelihoods of intralogous PPIs can predict the likelihood of PPIs in the hybrid. Again, the strengths of the interlogous PPIs were found to be correlated with intralogous PPIs in the hybrid (Figure 4D). Moreover, they are also correlated with the intralogous PPIs in parents (Figure 4E), corroborating the results from the large-scale proteomics data (Figure 3C and D). The intralogous PPIs in parents were found to be a stronger predictor than the intralogous PPIs in hybrid. This exception to the overall trend observed in the case of the large-scale proteomics data (Figure 3E) may be because of the sampling of the relatively small number of PPIs in the prefoldin network and of other methodological differences. Overall, the strengths of interlogous PPIs could be predicted from the intralogous PPIs in the hybrid as well as from those in parents (Figure 4F), confirming that chimeric PPIs occur as they are in parental species.

### Unexpected changes in PPIs in the hybrid

We found a high similarity of PPIs and protein abundance between the hybrid and its parents, as well as patterns consistent with the frequent assembly of chimeric complexes. In this section, we examine how likelihoods of PPIs in some protein complexes deviate from these expectations, keeping in mind our limited statistical power. We first examined the relative preference for either intralogous or interlogous PPIs in hybrid, which as noted earlier (Figure S10), suggests that both types of PPIs are equally likely to occur in hybrid and that many proteins show no clear preference for either type. Upon stratifying PPIs based on the relative preferences (z-score ≤ −1: prefer intralogous PPIs; z-score ≥ 1: prefer interlogous PPIs), we find that PPIs that do not have a clear preference occur between proteins that have less diverged orthologs (Figure S13). On the other hand, when PPIs have a higher relative preference for either intralogous or interlogous PPIs, we observe a higher divergence between orthologs (two-sided Mann-Whitney U test, P-value < 1e-4 for intralogous PPIs and P-value < 1e-3 for interlogous PPIs). This led us to examine whether the deviation of the interlogous PPIs from the predicted interlogous PPIs (from the parental intralogous PPIs (Figure 3E)) depends on the divergence of protein sequences between species. We found that orthologs that diverge more in sequence tend to deviate from their expected behaviour in hybrids (Figure 5A). Highly diverged orthologs are also more likely to participate in the interlogous PPIs with lower interaction scores than the intralogous PPIs in the hybrid and parents, suggesting occurrence of incompatibilities within interlogous PPIs (Figure S14). However, this effect might be limited to a small number of protein complexes where the interlogous PPIs have very low interaction scores and therefore, incompatibility is more pronounced. Additionally, because of the positive relationship between the protein abundance and the sequence conservation (Drummond et al. 2005), such incompatibilities would likely be dependent on the abundance of the proteins. Overall, although the signal is weak, these results suggest that proteins that diverge in sequence tend to assemble differently in hybrids, which may result from amino acid differences that affect binding directly or other processes that may affect the assembly of complexes in hybrids with specific functions that are less conserved at the protein level.

**Figure 5.**
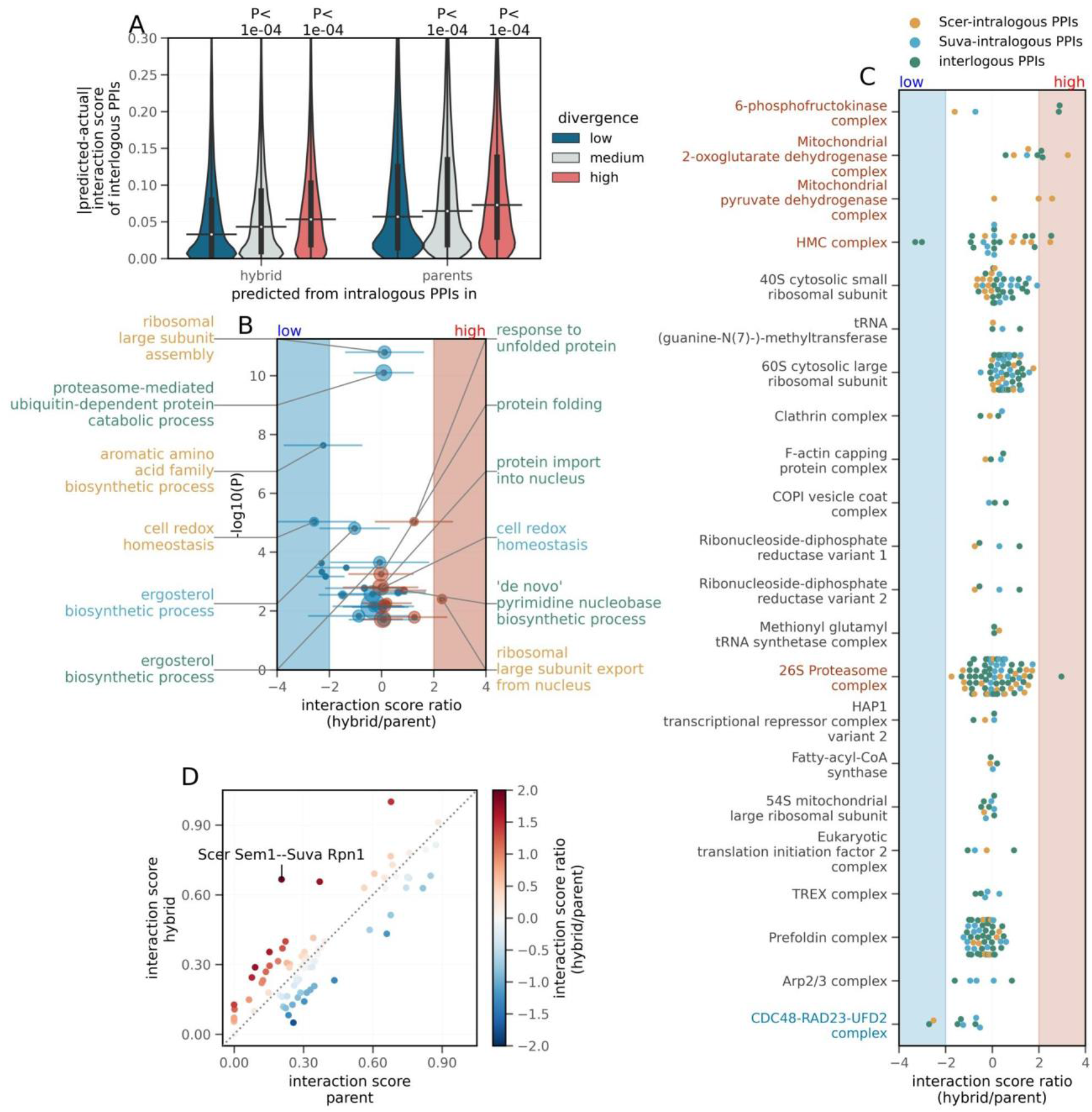
Differences of PPIs in hybrid compared to parental species. **A)** Relationship between the divergence of interlogous PPIs and the predictability of likelihood of the interlogous interactions. PPIs were categorised as between ‘low’, ‘medium’ or ‘high’ divergence interactors, based on the average of the divergence of interactor protein sequences, using 1st and 3rd quartiles as thresholds. Predictability of likelihoods of interlogous PPIs from intralogous PPIs in hybrid and parents is measured in terms of absolute difference between predicted and actual scores (y-axis). P-values from two-sided Mann-Whitney U tests are shown. On the violin plots, the medians of the distributions are shown by a horizontal black line and quartiles, by a vertical thick black line. **B)** Volcano plot showing the enrichment of GO terms from the Biological processes. Median value of z-score normalized hybrid/parent ratio of interaction scores. This is shown per set of PPIs corresponding to a gene set, on the x-axis. The -log10(P-value) for enrichment (hypergeometric test, FDR corrected by Benjamini & Hochberg method) is shown on the y-axis. Each point corresponds to sets of Scer-intralogous (yellow annotations), Suva-intralogous (blue annotations) or interlogous (green annotations) PPIs. For clarity, only the six most significantly low (on left) and significantly high (on right) enrichment gene sets are annotated. Size of the points is proportional to the number of genes per gene set detected in the proteomics data. Horizontal error bars indicate standard deviation of the ratio for the set of PPIs. The region shaded in blue indicates a significantly low ratio (z-score ≤ −2), while the region shaded in red indicates a significantly high ratio (z-score ≥ 2). **C)** Distributions of the z-score normalized hybrid/parent ratio of interaction scores per enriched protein complex are shown. Each point represents the ratio for a given PPI. The gene sets are sorted based on the median of ratio per gene set. The region shaded in blue indicates a significantly low ratio (z-score ≤ −2), while the region shaded in red indicates a significantly high ratio (z-score ≥ 2). Gene sets with significantly low or high ratios are colored in blue and red respectively. **D)** Comparison of the interaction scores of PPIs in hybrid (on y-axis) with those of parents (on x-axis), in the case of proteasome complex. Interaction scores of the intralogous PPIs in hybrid are compared with intralogous PPIs in parents, whereas interaction scores of the interlogous PPIs in hybrid are compared with interlogous PPIs in the theoretical hybrid. Color of the points indicates the ratio between the interaction scores in hybrid with respect to parents. The ratios are expressed as z-scores. The PPIs with a significantly different ratio (|z-score|≥2) are indicated on the plot.

In order to examine which biological functions may be particularly affected at the level of PPIs in hybrids, we looked at biological processes that have an enrichment of PPIs whose scores in hybrids are higher or lower than in the parents (see Methods). Remarkably, the gene set enrichment analysis revealed several significant changes in proteostasis related gene sets (Figure 5B). For example, gene sets related to protein folding and response to unfolded protein are enriched for PPIs with significantly high likelihoods of interlogous interactions in hybrid compared to those expected in the theoretical hybrid (P-value = 6e-04 and P-value = 9e-06 respectively, hypergeometric test, FDR corrected by Benjamini & Hochberg method). On the other hand, a gene set related to the proteasome mediated ubiquitin dependent protein catabolic process contains PPIs with significantly low relative likelihoods of interlogous interactions in hybrid (P-value = 8e-11). Also, gene sets related to export of the large subunit of the ribosome from nucleus contains PPIs with significantly high likelihoods of interaction in hybrid (P-value = 4e-03), whereas gene sets related to the assembly of large subunit of the ribosome contains few PPIs with significantly high and few PPIs with significantly low relative likelihoods (P-value = 7e-03 and P-value = 2e-11 respectively). Proteostasis related changes are also detected when we look at protein complexes (Figure S15A), where we find that the HMC protein complex (involved in protein folding) contains PPIs with significantly high relative likelihoods of Scer-intralogous interactions in hybrid (P-value = 1e-03), the proteasome (protein degradation) contains PPIs with significantly high relative likelihoods of interlogous interactions in hybrid (P-value = 5e-02), and the CDC48-RAD23-UFD2 complex (protein degradation) contains PPIs with significantly low relative likelihoods of interlogous and Scer-intralogous interactions in hybrid (P-value = 3e-04 and P-value = 2e-04 respectively). Additionally, gene set related to chaperone binding (molecular function) contains PPIs with significantly low likelihoods of Scer-intralogous interactions in hybrid compared to parents (P-value = 3e-03, shown in Figure S15B). Interestingly, an increase in the abundance of complexes associated with proteostasis has been observed in hybrids between species of Drosophila (Bamberger et al. 2018), suggesting that hybrids between species may be particularly prone to changes in proteostasis.

Apart from the proteostasis related genes, among biological processes, gene sets related to biosynthesis of aromatic amino acid, ergosterol and pyrimidine nucleobase also contain PPIs with significant changes in likelihoods in hybrid compared to parents. Some of the metabolic complexes that also show significantly stronger PPIs in hybrids are mitochondrial protein complexes that may reveal incompatibilities between the nuclear and mitochondrial genomes, which is known to take place among pairs of Saccharomyces species (Špírek et al. 2014). The interaction between the alpha and beta subunit of the 6-phosphofructokinase, a complex involved in glycolysis, shows stronger interlogous interactions in the hybrids, potentially affecting its function (Figure S16).

Across protein complexes, however, the distributions of the hybrid/parent ratio are centered at zero (Figure 5C), indicating that the likelihoods of PPIs in hybrid are similar to those in parents or those expected in a theoretical hybrid. Subtle exceptional changes in the likelihoods of PPIs are evident in the PPI networks of the significantly enriched protein complexes as shown in Figures S7 and S16 to S20. We illustrate such changes by focusing on the proteasome in the hybrid (Figure 5D). In this case, some interlogous PPIs seem favored over intralogous PPIs, suggesting existence of a certain degree of heterogeneity in the assembly of the complex. For instance, Scer Sem1p has a significantly higher likelihood of interaction with Rpn1p of Suva, compared to the expected interaction in the theoretical hybrid (z-score≥2). The potential changes in the assembly of the proteasome raises interesting links with proteostasis as the proteasome is the main complex involved in protein degradation. Whether the particular changes in the assembly of the proteasome is one of the causes or the consequences of the perturbation of proteostasis in hybrids will require further investigation.

## Conclusion

Given the functional importance of protein complexes in the cell, variation in their organization may play a central role in the cellular phenotypes of hybrids. In this study, we investigated the molecular basis differentiating hybrids from parents by looking directly at their PPIs. Using the proteomics approach of SEC-PCP-SILAC (Havugimana et al. 2012; Kristensen et al. 2012) (Figure 1B), we obtained likelihoods of physical associations between proteins for *Saccharomyces cerevisiae* × *Saccharomyces uvarum* F1 diploid hybrid and their parents. We investigated the assembly of a few dozen protein complexes in the hybrid in a comparative manner. In general, accurate identification of interacting protein pairs in datasets resulting from co-migration experiments is known to be challenging (Skinnider et al. 2018). The relatively small number of proteins identified for each species was one of the limiting factors of the study. Because of the technical constraints related to the proteomics method, in order to confidently ascribe the parental origin of the proteins, we resorted to one of the most diverged pairs of parental species in the Saccharomyces phylogeny and could only utilize the uniquely aligned peptides for resolving the species origin for the proteins in hybrid. However, a lack of unduly bias for highly divergent proteins among the detected proteins suggests that protein sequence divergence of the orthologs was not the only factor limiting the number of proteins in the dataset. The resulting subsampling of the proteome due to the technical constraints related to the proteomics approach we adopted limited the extent of whole proteome-wide extrapolations from our study. For a broader coverage of the proteome, among other things, the use of additional labeled amino acids could be advantageous. Further investigation on the dependence of the frequency of chimeric complexes on divergence between parental species might also require direct protein interaction assays.

The overall equally likely intralogous and interlogous PPIs in hybrids suggest frequent assembly of chimeric protein complexes in hybrid and thus that the subunits of complexes are generally replaceable at this phylogenetic distance, consistent with the results of (Leducq et al. 2012). However, our experimental analysis is biased towards large, abundant and thus possibly more essential and stable complexes, which might be less sensitive to protein divergence for their assembly. Yeast F1 hybrids rarely show reduction of fitness compared to parents in terms of growth rate (Bernardes et al. 2017; Charron & Landry 2017) defects of sporulation and spore survival (Lee et al. 2008; Greig 2009; Charron et al. 2014). It may therefore be expected that protein complexes involved in the core cellular machinery such as the one studied here assemble correctly in hybrids. Additionally, the analysis is based on the well-annotated protein complexes of Scer, with the assumption that the composition of protein complexes is mostly similar between Scer and Suva. Though, high overall similarity in likelihoods of the PPIs supports the assumption, it might have limited the detection of subtle differences in the assembly of protein complexes. We also did not thoroughly explore the possibility of novel complexes in hybrids because we had limited technical power for such discovery. Another element to consider is that the strains were grown in synthetic media and differences between hybrids and parents could be manifested in specific conditions such as during stress responses, which may trigger the activity of proteins and pathways that have diverged between species.

In spite of this large-scale conservation of PPIs in hybrids, we identified some biological functions that seem to be altered specifically in hybrids, including the ones related to proteostasis and metabolism. This could be an effect of or a response to misregulation of protein proteostasis in hybrids, as has been observed for many other hybrid molecular traits (Landry et al. 2007; Bar-Zvi et al. 2017). Protein complexes involved in proteostasis such as the proteasome could be altered in hybrids, resulting in perturbation of the associated functions. Another possibility is that there could be some imbalance among protein subunits of complexes in the hybrids or marginal incompatibilities among subunits, which would put an additional burden on the protein quality control machinery because the regulation of the stoichiometry of large complexes often occurs through excess protein degradation (Taggart & Li 2018). This hypothesis is supported by a weaker dosage balance for interlogous interacting protein pairs in the hybrid compared to intralogous ones in parents. Proteostasis has also been noticed as being enhanced in hybrids of Drosophila (Bamberger et al. 2018) raising the interesting possibility that protein-protein or gene expression incompatibilities in hybrids are of weak effects but distributed among many proteins, imposing a general proteomic stress. In conclusion, protein complexes appear to be generally robust to protein divergence between species but small differences in assembly or abundance could accumulate to a point where they perturb the overall protein physiology of the cell. An interesting avenue of research would be to examine how complexes accommodate the evolution of the hybrid genomes with time, for instance through recombination and loss of heterozygosity (Zhang et al. 2020), which may further enhance the imbalance of complexes.

## Material and Methods

### Generation of yeast hybrids for the SEC-PCP-SILAC experiment

The hybrids generated in this study are described in Tables S1A and S1B along with the replicates that were performed (Table S1C). Parental strains used to generate the hybrids are derived from the reference strains S288C (*Saccharomyces cerevisiae*) and CBS7001 (*Saccharomyces uvarum*). Using a microneedle (SporePlay micromanipulator, Singer Instruments), a single haploid cell of the first strain was put in physical contact with a single haploid cell of the second strain on a YPD plate (Table S2). After three days of growth at 30 °C, flow cytometry (Millipore Guava easyCyte Flow Cytometry System, MilliporeSigma) was used to determine the relative ploidy of colonies, as reported in (Gerstein et al. 2006). Diploid hybrid clones were then validated by PCR on the mating type locus (Huxley et al. 1990). Colony PCRs were performed on single colonies as follows: a small amount of fresh colonies was resuspended in 40 μl of NaOH 20 mM and incubated for 20 min at 95 °C for cell lysis. Cells were centrifuged for 5 min at 4000 rpm. For each PCR reaction, the mixture contained 2.5 μl of 10X BioShop^®^ Buffer, 2 μl of supernatant of the lysed cells, 1.5 μl of MgCl2 25 mM, 0.5 μl of dNTP mix 10 mM, 0.5 μl of each primer at 10 μM and 0.15 μl of Taq DNA Polymerase (5 U/μl, BioShop Canada Inc.), in a final volume of 25 μl. PCR reactions were carried out in a thermocycler (MasterCycler ProS, Eppendorf©) with the following steps: 5 min at 95 °C; 35 cycles of 30 s at 94 °C, 30 s at 55 °C and 1 min at 72 °C; and a final extension of 3 min at 72 °C. PCR products were then size-verified on agarose gel. Primers used for verification of mating type are described in Table S3. After selection of diploid cells, the strains involved in the crosses were validated by analysis of the restriction profiles obtained after DNA digestion with AccI enzyme: quick DNA extraction was performed on single diploid yeast colonies (Lõoke et al. 2011) followed by PCR amplification using universal primers for Saccharomyces species in the POP2 gene (Table S3). PCR reaction mixture was prepared as described above and incubated following these steps: 5 min at 95 °C; 35 cycles of 30 s at 95 °C, 30 s at 52 °C and 2 min at 72 °C; and a final extension of 10 min at 72 °C. Each PCR amplification product was digested overnight at 37 °C with 2 μl of CutSmart^®^ uffer (New England Biolabs) and 0.2 μl of AccI enzyme (10 U/mL, New England Biolabs). Restriction profiles specific to each species (Scer or Suva) were identified on agarose gel.

### SILAC labeling

SILAC labeling was performed as described by (Fröhlich et al. 2013). All yeast strains used in this study were auxotrophic for lysine. SC (Synthetic Complete) -lys medium (Table S2) was prepared and enriched by adding 30 mg/L of the following isotopes: (1) L-Lysine (Sigma-Aldrich) for “light” (L) labeled cells, (2) D4 L-Lysine (Cambridge Isotope Laboratories) for “medium” (M) labeling and (3) ^13^C6 ^15^N2 L-Lysine (Cambridge Isotope Laboratories) for “heavy” (H) labeling. Cells were pre-cultured overnight in 5 mL of medium containing L, M or H lysine at 25 °C. Two 50 mL cultures (L samples corresponding to parent Scer (50 mL) and parent Suva (50 mL)) and two 100 mL cultures (M and H samples) of medium with corresponding isotope were inoculated from the pre-cultures to A_600_ = 0.001. For comparison between a hybrid and a parent, M sample corresponded to one parent, either Scer or Suva (100 mL), and H sample corresponded to one biological hybrid (100 mL). For comparison between parents, M sample corresponded to parent Scer (100 mL) and H sample corresponded to the other parent Suva (100 mL). Cells were grown to a final A_600_ = 0.7 corresponding to more than ten doublings. The labeling efficiency was found to be between 85-88% (ratio of number of proteins with M or H modifications on total number of proteins). Summary of the labeled samples can be found in Figure S2.

### Cell lysis

After SILAC labeling, the two parental L samples were pooled to obtain the same final volume than M and H samples (100 mL). Cells were harvested via centrifugation and washed twice with cold Tris-Buffered Saline (TBS) solution (50 mM Tris and 150 mM NaCl at pH 7.5). Cells were resuspended in 500 μL of SEC lysis buffer (50 mM Tris, 50 mM NaOAc and 50 mM KCl at pH 7.2) including Halt™ Protease and Phosphatase Inhibitor Cocktail (100X) (ThermoFisher Scientific) and then quickly frozen as droplets into liquid nitrogen. The cells were lysed by grinding with a cold mortar and pestle in liquid nitrogen. After lysis, 2.5 ml of SEC lysis buffer with protease inhibitors were added to each lysate. To enrich soluble and cytosolic complexes, the obtained volume was clarified by ultracentrifugation (100 000 rcf for 15 min at 4 °C). A protein concentration assay was performed on supernatant (Pierce BCA Protein Assay Kit, Thermo Fisher Scientific) to inject the same amount of proteins for each sample (L, M, H) in the Size-Exclusion Chromatography (SEC) column. The minimum total amount of proteins injected into the SEC column was 700 μg and the maximum amount was 1000 μg. Proteins and protein complexes were then extracted and the efficiency of protein extraction was validated with SDS-PAGE and native gels for both hybrids and parents (Figure S21A to C).

### Size-Exclusion Chromatography (SEC)

The SEC samples were divided into two technical replicates. As described by (Kristensen et al. 2012), to reduce the volume and to enrich for high-molecular weight complexes, samples were concentrated using ultrafiltration (100,000 MWCO, Sartorius Stedim). The L sample was concentrated to 200 μL (two technical replicates) while M/H samples were combined just before loading into the SEC column (100 μL + 100 μL) (two technical replicates). Note that technical replicates started from the same cell cultures but samples were divided in two just before injection onto the SEC machine. Biological replicates are experiments starting from independent cultures and different hybrid crosses (Figure S2). The fractions from the L samples served as an internal standard and were separated by SEC independently from the M/H samples. Samples were loaded into a 300 x 7.8 nm BioSep4000 Column (Phenomenex) and separated into 80 fractions by a 1200 Series semi-preparative HPLC (Agilent Technologies) at a flow rate of 0.5 mL/min at 8 °C. The collected volume of the first twenty fractions was 250 μL per fraction and decreased to 125 μL per fraction for fractions 21-80. After protein extraction, SEC separation was performed on the L sample on one side, and M/H samples on the other side. Elution profiles were used as indicators of proper protein complexes separation without aggregates or protein complex dissociation (Figure S21D). In total, 20 samples were separated by the SEC. The information regarding the technical replicates and the samples is provided in Table S1C.

### Protein digestion

Individual SEC-PCP-SILAC samples were prepared for digestion as described in (Scott et al. 2015). The first five fractions were skipped as they likely contain the void volume and protein aggregates. The last fractions (66 to 80) were also skipped to keep only the most abundant cytosolic complexes and to eliminate smaller-sized proteins (most probably non-interacting proteins). A total of 60 fractions (6 to 65) were thus used in the following steps. Urea (6 M) and thiourea (2 M) were added to each M/H fraction. To generate the SEC-PCP-SILAC reference, fractions 6 to 65 of the L SEC separated samples were pooled together and added to each of the M/H fractions at a volume of 1:1. Ammonium bicarbonate (50 mM) was added to the fractions to stabilize the pH. Disulfide reduction was performed by incubating each fraction with 1 ug dithiothreitol (DTT) for 30 minutes at room temperature. Samples were then alkylated with 5 μg iodoacetamide (IAA) in the dark for 20 minutes at room temperature. LysC was added at a ratio of 1:50 and samples were incubated again overnight at 37 °C. Samples were acidified to a pH < 2.5 with 20% trifluoroacetic acid (TFA). Peptides were then purified using self-made stop-and-go-extraction tips (StageTips, (Rappsilber et al. 2003)) made with C18 Empore material packed into 200 μL pipette tips. Stage Tips were first conditioned with 100 μL of methanol and equilibrated with 1% TFA. Peptides were loaded into the column and then washed twice by adding 100 μL of 0.1% formic acid followed by centrifugation. Peptides were finally eluted with 100 μL of 0.1% formic acid and 80% acetonitrile. Samples were dried down using a vacuum concentrator and stored at 4 °C.

The SEC-PCP-SILAC steps described in this protocol were adapted from (Kristensen et al. 2012).

### Mass spectrometry

Prior to the mass spectrometry analysis, samples were resuspended in 30 uL (fractions 6 to 20) or 15 uL (fractions 21 to 65) of 0.1% formic acid. Peptides were analysed using a quadrupole-time of flight mass spectrometer (Impact II, Bruker Daltonics) on-line coupled to an Easy nano LC 1000 HPLC (ThermoFisher Scientific) using a Captive spray nanospray ionization source (Bruker Daltonics) including a 2-cm-long, 100-μm-inner diameter fused silica fritted trap column, a 40-cm-long (average), 75-μm-inner diameter fused silica analytical column with an integrated spray tip (6-8-μm-diameter opening, pulled on a P-2000 laser puller, Sutter Instruments). The trap column is packed with 5 μm Aqua C18 beads (Phenomenex) while the analytical column is packed with 1.9 μm-diameter Reprosil-Pur C18-AQ beads (Dr. Maisch, www.Dr-Maisch.com). The analytical column was held at 50 °C by an in-house constructed column heater. Buffer A consisted of 0.1% formic acid in water, and buffer B consisted of 0.1% formic acid in acetonitrile. Peptides were separated from 0% to 40% Buffer B in 90 minutes, then the column was washed with 100% Buffer B for 20 minutes before re-equilibration with Buffer A.

The Impact II was set to acquire in a data-dependent auto-MS/MS mode with inactive focus fragmenting the 20 most abundant ions (one at the time at 18 Hz rate) after each full-range scan from m/z 200 Th to m/z 2000 Th (at 5 Hz rate). The isolation window for MS/MS was 2 to 3 Th depending on parent ion mass to charge ratio and the collision energy ranged from 23 to 65 eV depending on ion mass and charge. Parent ions were then excluded from MS/MS for the next 0.4 min and reconsidered if their intensity increased more than 5 times. Singly charged ions were excluded since in ESI mode peptides usually carry multiple charges. Strict active exclusion was applied. The nano ESI source was operated at 1700 V capillary voltage, 0.20 Bar nano buster pressure, 3 L/min drying gas and 150 °C drying temperature. The mass spectrometry data are available through PRIDE (Perez-Riverol et al. 2019) at https://www.ebi.ac.uk/pride/archive/projects/PXD010136.

### Database searching and quantification

Tandem mass spectra were extracted from the data files using MaxQuant version 1.6.0.1 (Cox & Mann 2008) and were searched against protein sequences from Scer and Suva retrieved from Saccharomyces Sensu Stricto online resources (http://www.saccharomycessensustricto.org/) (Scannell et al. 2011) plus common contaminants and reverse database for false discovery rate (FDR) filtering. Peptide and protein identification was performed using Maxquant (Tyanova et al. 2016). MaxQuant was used to identify proteins in our samples with the following parameters: carbamidomethylation of cysteine as a fixed modification; oxidation of methionine, acetylation of protein N-terminal and SILAC labeling as variable modifications; Lysine/K cleavage with a maximum of two missed cleavages, 0.006 Da precursor mass error tolerance and 40 ppm fragment ion mass tolerance and requantify option was enabled. The data was filtered for 1% FDR at both peptide and protein level. The search results and the protein databases are available at https://www.ebi.ac.uk/pride/archive/projects/PXD010136.

### Preprocessing of proteomics raw data

From the ‘proteinGroups’ output of MaxQuant, proteins (column name: ‘Protein Groups’) with commonly occurring contaminant (column name: ‘Potential contaminant’), proteins identified only by a modification site (column name: ‘Only identified by site’), and other spurious protein hits (column name: ‘Reverse’) were discarded. Additionally, protein hits with Andromeda score (column name: ‘Score’) of less than 0.05 quantile threshold were discarded, to retain only high quality hits. The filtered protein hits were annotated by gene name/id based on the proteome reference, and by species of origin based on the SILAC labeling (Figure S2). Only uniquely aligned peptides were considered while assigning the species of origin, avoiding ambiguity in the data. The L labeling served as a reference for the calculation of the ratios of intensities i.e. M/L and H/L. The ratios of intensities along the elution fractions constituted an elution profile for a given peptide. Such peptide-wise data and replicates were aggregated by taking the mean of the ratios of intensities per elution profile, resulting in protein-wise elution profiles. During this procedure, aggregation of replicates helped in reducing the sparsity of the data. The data for the parental strains labeled with M and H isotopes were considered as biological replicates. Processed elution profiles of the proteins are included in Data S1.

### Estimation of interaction scores from proteomics data

The peak heights in the preprocessed protein-wise elution profiles were first rescaled between 0 and 1, so that minimum peak height is 0 and maximum peak height is 1. The rescaled elution profiles were used to calculate pairwise similarity (in ‘all versus all’ manner) using Dynamic Time Warping (DTW), implemented through dtaidistance (Wannesm et al. 2019). For the optimization of the window size parameter of the DTW, using the database reported interactions for Scer as a reference, we scanned a continuous range of window sizes (Figure S3). Reference PPIs were obtained from STRING (release version: 11.0, accessed on 05-11-2019) (Szklarczyk et al. 2019) and HitPredict (accessed on 08-11-2019) (López et al. 2015) databases. Based on the distance scores obtained from the DTW, an upper threshold was selected which marks the ‘no interaction’ range, with most accuracy. PPIs with distance scores higher than the upper threshold were assigned an interaction score of 0, indicating no interaction. If the distance score was less than the upper threshold, the interaction score was scaled in a way that the highest interaction score was equal to the lowest distance score and vice versa (interaction score = 1 - (distance score/upper threshold of distance score)). After optimizing the window size and the upper threshold of the distance score using parent Scer data, the interaction scores for Suva and hybrid were estimated using the optimized settings. Estimated interaction scores of the proteins are included in Data S2.

### Estimation of protein abundance from proteomics data

From the ‘proteinGroups’ output of MaxQuant, the ‘Peptide counts (all)’ column was used to estimate the protein abundances. The sum of the counts for all peptides of the same protein was used as protein abundance. In order to remove low spurious peptide counts, total counts of less than three were discarded. The protein-wise aggregated abundance was transformed with pseudo-log (base 10 with pseudocount of 0.5). Finally, the protein abundances were quantile normalised across species. Note that in the case of the hybrid, protein abundances for Scer as well as Suva proteins were estimated.

As shown in Figure 2B, as a validation of the protein abundances estimated from the proteomics data, we compared protein abundance obtained from the proteomics for parental Scer species with the reference protein abundance obtained from proteomics source of PeptideAtlas (Desiere et al. 2006) (March 2013, filename: 4932-PA_201303.txt, accessed on 23-05-2018) available on PAXdb database (Wang et al. 2015).

Dosage balance was measured as `1-((|p1-p2|)/(p1+p2)),` where `p1` and `p2` are protein abundances of two proteins. Protein abundance and the dosage balance scores are included in Data S3.

### Comparative analysis of the interaction scores between hybrid and parent species

The difference in the interaction scores of hybrid compared to parents was estimated in terms of a ratio (on log2 scale). The ratios were z-score normalised. The resulting distribution of z-score normalised ratios was used to stratify the PPIs with z-score greater than 2 as PPIs with significantly ‘high’ relative likelihoods in hybrid and those with z-score less than −2 as with significantly ‘low’ relative likelihoods in hybrid. The ratios of interaction score are included in Data S5.

### Gene set enrichment analysis

Reference gene sets used in the enrichment analysis include sets of protein complexes that were obtained from complex portal (Meldal et al. 2019), and GO (Gene Ontology) terms obtained from QuickGO (Binns et al. 2009). In order to avoid redundancy in the set of large protein complexes, if a given protein complex possesses different variants (e.g. proteasome, ID: CPX-2262), only the variant carrying the largest number of interactors was considered in the analysis. Among GO terms, gene sets that qualify as ‘part_of’, ‘involved_in’ or ‘enables’ from Molecular function (F), Biological Process (P) and cellular component (C) were included in the dataset. Only gene sets assigned by GO_Central (Binns et al. 2009; The Gene Ontology Consortium & The Gene Ontology Consortium 2019) or SGD (Griffith & Griffith 2004) were retrieved from QuickGO. Link to access the GO terms used in the study: https://www.ebi.ac.uk/QuickGO/annotations?qualifier=part_of,involved_in,enables&assignedBy=GO_Central,SGD&reference=PMID&taxonId=559292&taxonUsage=descendants&geneProductSubset=Swiss-Prot&proteome=gcrpCan,gcrpIso,complete&geneProductType=protein&withFrom=GO,SGD

As a test gene sets, the sets of PPIs with significantly ‘high’ and ‘low’ interaction scores obtained from the comparative analysis of the hybrid with parents were used. The significance of the overlap between the reference and test gene sets was tested with hypergeometric test and it was corrected for false discovery rate (FDR) using Benjamini-Hochberg procedure. The results of the gene set enrichment analysis are included in Data S5.

### DHFR-PCA screening of the prefoldin complex PPIs

#### Strain construction

The DHFR-PCA method was applied to detect PPIs between proteins forming the prefoldin complex in diploid cells. Multiple steps were needed to construct these strains (Tables S1A and S1C).

#### Single-tagged haploid strains

First, Scer haploid MATa (BY4741) and MAT**α** (BY4742) strains were retrieved from the Yeast Protein Interactome Collection (Tarassov et al. 2008) (except for strain with PFD1 gene tagged with DHFR F[3] which was reconstructed as specified below for Suva). Second, Suva haploid MATa (MG032) and MAT**α** (MG031) strains were constructed as follows. DHFR fragments and associated resistance modules were amplified from plasmids pAG25-linker-F[1,2]-ADHterm (NAT resistance marker) and pAG32-linker-F[3]-ADHterm (HPH resistance marker) (Tarassov et al. 2008) using oligonucleotides described in Table S3. PCR mixture contained 15 ng of plasmid, 5 μL of 5X Buffer with Mg2+, 0.75 μL of 10 mM dNTPs, 3 μL of each primer at 10 μM and 0.5 μL of 1 U/μL Kapa HiFi HotStart DNA polymerase (Kapa Biosystems, Inc, A Roche Company) for a total volume of 25 μL. PCR was performed with the following cycling protocol: initial denaturation (5 min, 95 °C), 32 cycles of 1) denaturation (20 s, 98 °C), 2) annealing (15 s, 64,4 °C) and 3) extension (1 min, 72 °C) and one cycle for final extension (5 min, 72 °C). PCR products were then concentrated using an OligoPrep OP120 SpeedVac Concentrator (Savant). Competent cells collected during exponential growth (A_600_ = 0.7) were transformed with either DHFR F[1,2] (MATa cells) or DHFR F[3] (MAT**α** cells) modules as described in (Tarassov et al. 2008) with the following modifications: heat shock was performed for 20-30 min after adding 5 μL of Dimethyl Sulfoxide (DMSO), followed by recovery in YPD at 25 °C for five hours. Cells were plated onto selective NAT (DHFR F[1,2]) or HYG (DHFR F[3]) media (Table S2) and incubated for five days at 25 °C. Correct genome integration of DHFR fragment module was validated by colony PCR using primers described in Table S3. Colony PCRs were performed as mentioned previously (mating type verification PCR), with the only difference that we used 2,5 ul of supernatant of lysed cells for each PCR reaction. The product of the PCR reaction was finally Sanger sequenced with O1-50 primer ensuring that no insertion, deletion, or non-synonymous mutation occured at the junction between the gene and the DHFR fragment.

#### Double-tagged haploid strains

At this point, we had all the haploid strains with one of the prefoldin complex genes tagged with either DHFR F[1,2] or DHFR F[3] (Table S1A). These strains would be used to test PPIs in parents and interlogous PPIs in hybrids. But to test for parental PPIs in hybrids, we needed single haploid strains tagged for both prefoldin genes for which we want to test the interaction, one tagged with DHFR F[1,2] and the other with DHFR F[3] (see Figure S11).

To achieve that, for Scer strains, we crossed previously described MATa and MAT**α** haploid tagged strains of interest to obtain all desired combinations. Briefly, cells of opposite mating types were combined into 3 mL of YPD. Cells were incubated overnight at 30 °C and diploid selection was performed on NAT+HygB (Table S2). Cells were then transferred onto enriched sporulation medium and incubated for at least one week at room temperature. Following ascus digestion with 200 μg/mL zymolyase 20T (BioShop Canada Inc.), sporulated cultures were put on solid YPD medium. Tetrads were then dissected with a microneedle (SporePlay micromanipulator, Singer Instruments) to isolate single haploid spores. Mating type of these spores was PCR identified following Huxley et al. procedure described above and spores were replicated on SC -met and SC -lys (Table S2) to determinate auxotrophies. Selected strains for the following experiment are identified in Table S1D.

For Suva strains, we used an alternative approach. We transformed MG032 haploid strains already tagged for one gene with DHFR F[1,2] directly with specific DHFR F[3] modules amplified from a pAG32-linker-F[3]-ADHterm plasmid in which the TEF terminator for the antibiotic resistance was changed for a CYC terminator (pAG32-DHFR[3]-HPHNT1) to avoid unwanted recombination between the resistance markers. We performed the same steps as described above with the following changes: reverse primers used for plasmid amplification were different and are described in Table S3 while forward primers remained the same. After transformations, cells were plated on NAT+HygB. The following steps remained similar, including validation of transformations by colony PCRs and sequencing.

For positive controls, we transformed Scer BY4741 MATa and BY4742 MAT**α** strains with plasmids p41-ZL-DHFR[1,2] and p41-ZL-DHFR[3]. As described by (Leducq et al. 2012), these plasmids express interacting leucine zipper moieties that strongly dimerize and that thus lead to a strong signal in DHFR-PCA. For negative controls, we used plasmids expressing the linkers and DHFR fragments alone, p41-L-DHFR[1,2] and p41-L-DHFR[3], that show no signal in DHFR-PCA.

### Screening of PPIs in the prefoldin complex

We performed all possible crosses among the constructed strains (see above and Table S1D) for a total of 15 different tested PPIs and 180 independent crosses. MATa and MAT**α** strains were combined from solid medium into a 96-well deepwell plate with 1 mL of YPD per well. About the same amount of cells for each strain were transferred into each well. Plates were incubated overnight at 25 °C. The next day, cells were resuspended and 6 μl of each cross was deposited on solid SC medium lacking the proper amino acids to ensure diploid selection (Tables S1D and S2). Plates were incubated for at least two days at 25 °C. After diploid selection, cells were transferred again into a 96-well deepwell plate with 1 mL of liquid SC medium lacking the proper amino acids and put at 25 °C overnight. The following days, cells were printed and then rearrayed on YPD plates in a way to have in a 1536 format a minimum of six replicates per diploid strains and to include a double border of a control PPI of medium strength (LSM8-DHFR F[1,2]/CDC39-DHFR F[3]). To perform DHFR-PCA, cells were in the end transferred on MTX and DMSO (ctl) media (Table S2) for two successive rounds of a four-day incubation at 25 °C. Starting from the rearraying step, all the following steps were done using robotically manipulated pin tools (BM5-SC1, S&P Robotics Inc).

### Estimation of interaction scores from DHFR-PCA experiment data

Images of agar media plates used for the DHFR-PCA experiment were taken each day of the two selection rounds with an EOS Rebel T5i camera (Canon). We used images taken after four days of growth on the second selection round for analysis of both MTX and DMSO plates. Images were analysed using gitter (R package version 1.1.1; (Wagih & Parts 2014)) to quantify colony sizes (Data S6) by defining a square around the colony center and measuring the foreground pixel intensity minus the background pixel intensity. For the estimation of the interaction scores (Data S6), first, log2-scaled ratios of sizes of the colonies on MTX with respect to sizes of the colonies on the DMSO plates were calculated. The ratios calculated from crosses representing the same interaction types (as shown in Figure S11) were averaged. All the ratios were rescaled between 0 (no interaction) and 1 (strong interaction). The rescaled ratios are referred to as interaction scores in the text.

## Availability of code and data

Supplementary data files are available on figshare at https://doi.org/10.6084/m9.figshare.12428975.v1. Raw proteomics data is available on PRIDE data repository at https://www.ebi.ac.uk/pride/archive/projects/PXD010136. Computer code to reproduce all the analyses in this manuscript is available at https://github.com/Landrylab/dandage_berger_2020.

## Acknowledgements

We thank the members of the Landrylab for discussions. We thank Anna Fijarczyk, Angel F. Cisneros, Johan Hallin and Diana I. Ascencio for comments on the manuscript. CMB thanks the teams of LJF and Thibault Mayor for their help in the project and access to equipment. RD is funded by Fonds de recherche du Québec-Santé (FRQS) Programme Postdoctoral and HFSP (RGP0034/2018) grant to CRL. CMB was funded by PROTEO via a FQRNT/PROTEO International Internship Program. This research was supported by the NSERC discovery grant to CRL and by Genome Canada and Genome British Columbia to LJF (Project 214PRO). CRL holds the Canada Research Chair in Canada Research Chair in Cellular Systems and Synthetic Biology. The mass spectrometry infrastructure used here was supported by the Canada Foundation for Innovation, the British Columbia Knowledge Development Fund and the BC Proteomics Network.

## Author’s contributions

CRL, IGA and LJF designed research. CMB, RGS, IGA and KMM performed experiments. RD and CMB performed the analyses. RD, IGA and CRL wrote the paper with input from CMB.

## Declaration of interest

None to declare.

## Notes

### Competing Interest Statement

The authors have declared no competing interest.

