## Supporting Information for "Frequent assembly of chimeric complexes in the protein interaction network of an interspecies yeast hybrid"

### *Supplementary figures*

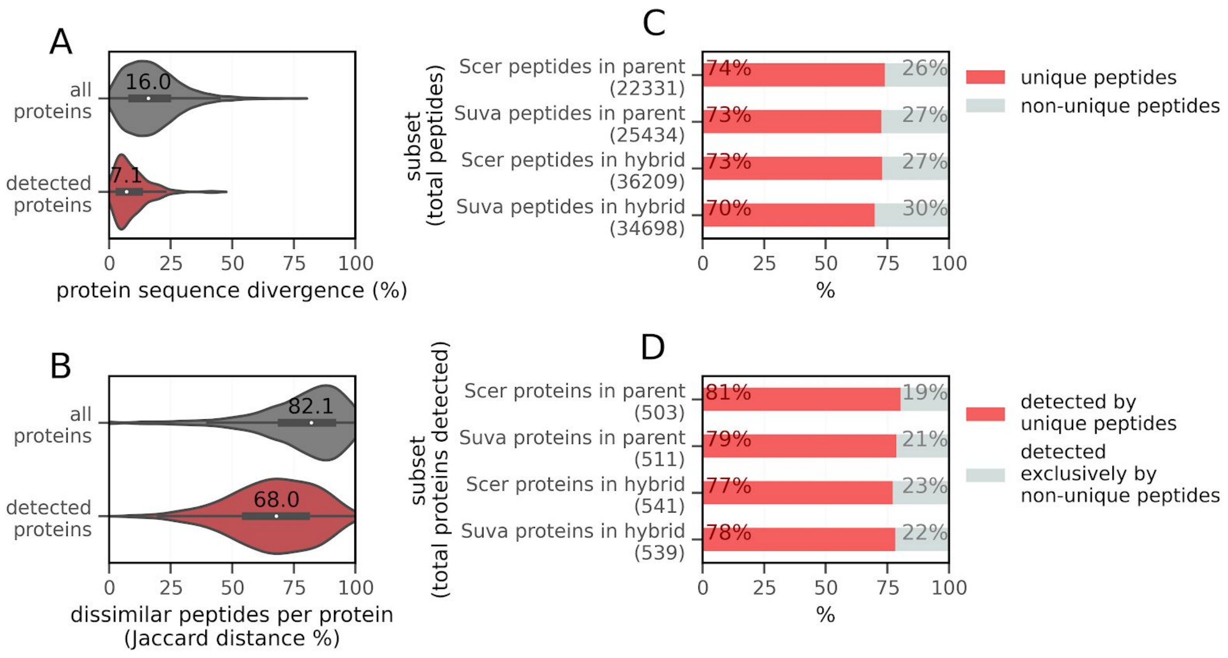


**Figure S1. *Saccharomyces cerevisiae* and *Saccharomyces uvarum* parental strains are divergent enough for the identification of proteins from SEC-PCP-SILAC.**

**A)** Ortholog divergence between Scer and Suva. Orthologs were aligned using Muscle (Edgar 2004) and the level of pairwise amino acid divergence for each pair of ortholog was determined.

**B)** Peptide divergence between orthologous Scer and Suva proteins. Simulations of the protein digestion by LysC enzyme (which cuts after each lysine) were carried out using pyOpenMS (Röst et al. 2014). Peptides from the same protein were then compared between Scer and Suva in terms of Jaccard distance. Peptide divergence is shown for proteins detected in the proteomics experiment and for all parental species proteins.

**C)** Stacked bar plot showing the percentage of uniquely aligned peptides, out of the total number of peptides detected by mass spectrometry. Only the filtered set of uniquely aligned peptides was used in the analysis.

**D)** Stacked bar plot showing the percentage of proteins detected by uniquely aligned peptides, out of the total number of proteins detected by mass spectrometry. Only the filtered set of proteins detected by the uniquely aligned peptides was used in the subsequent analysis.

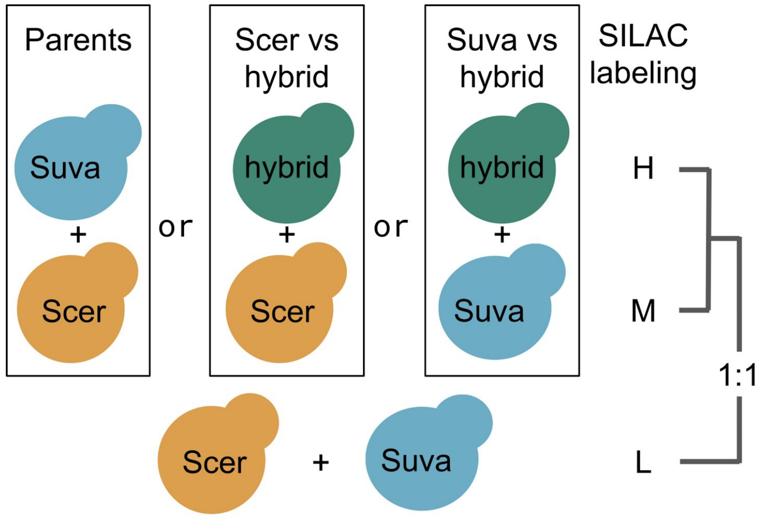


**Figure S2. SILAC labeling.**
Parental and hybrid strains were labeled with unique isotopes of lysine. Medium (M) and heavy (H) isotopes are used for interspecies comparison, while light isotope (L) is used to generate reference chromatograms. After SEC, the L fractions are pooled and added in equal amounts into M/H fractions.

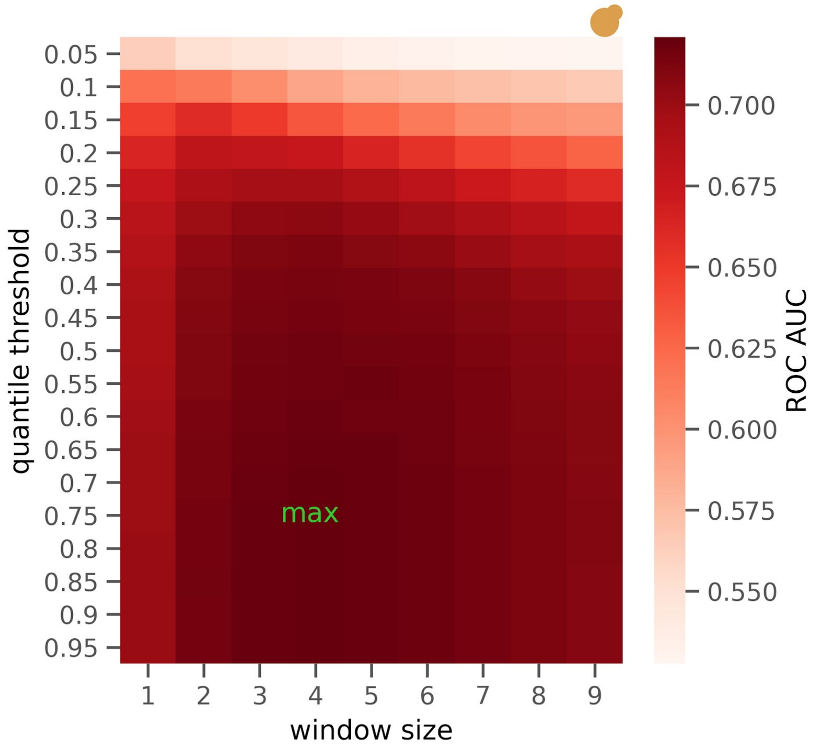


**Figure S3. Optimization of parameters to capture interactions from proteomics data.**

Range of window sizes in terms of number of elutions used for DTW (x-axis) and range of quantile thresholds (y-axis) were assessed to find the combination that provides maximum accuracy (denoted as 'max' on the heatmap). The accuracy was measured in terms of Area Under the Curve of Receiver Operating Characteristic i.e. ROC AUC. Note that interactions used in this analysis correspond to the Scer parent. See Methods section for the details of the optimization strategy. The logo of Scer indicates that this analysis was carried out only with the parental Scer data.

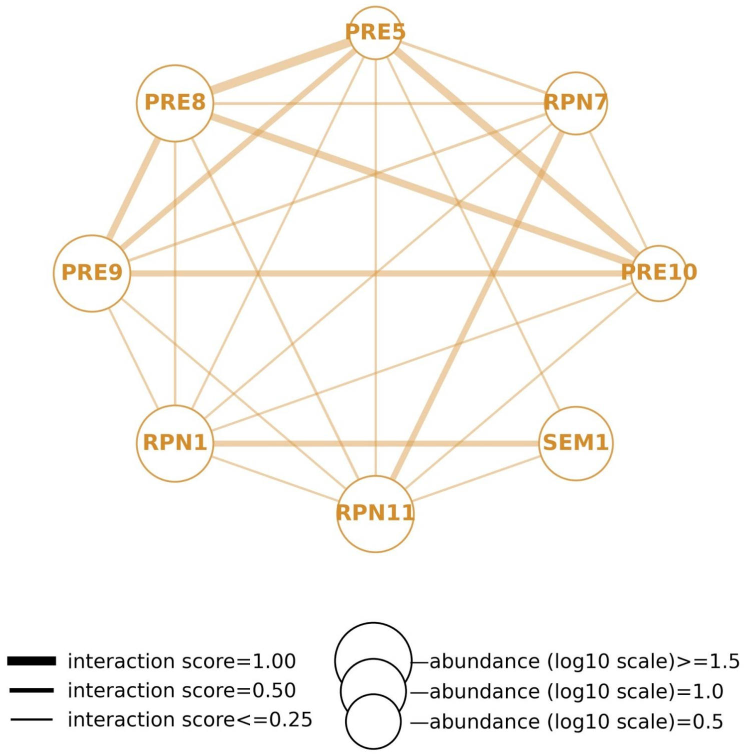


**Figure S4. PPI network within Scer parent proteasome, as monitored by SEC-PCP-SILAC.**

The sizes of the nodes are in scale with protein abundance and the width of the edges are in scale with interaction scores.

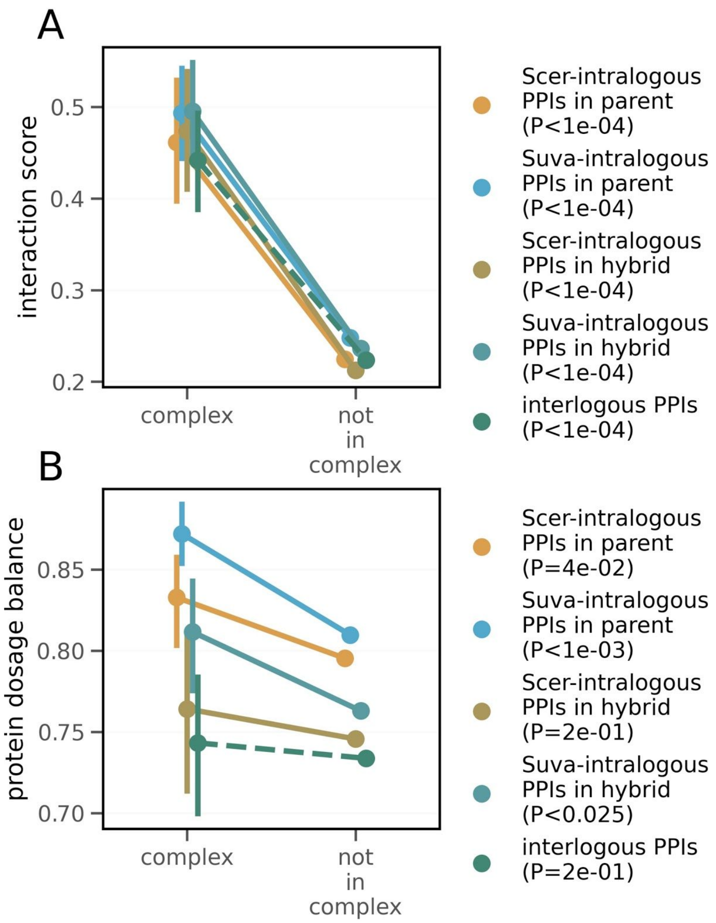


**Figure S5. Comparison of interaction scores (A) and dosage balance (B) between interactors within complexes with that of interactors that are not contained in a protein complex.**

**A)** Comparison of the interaction score for PPIs within protein complexes with those not contained in any protein complex.

**B)** Comparison of the dosage balance for PPIs within protein complexes with those not contained in any protein complex.

Bars denote a 95 % confidence interval.

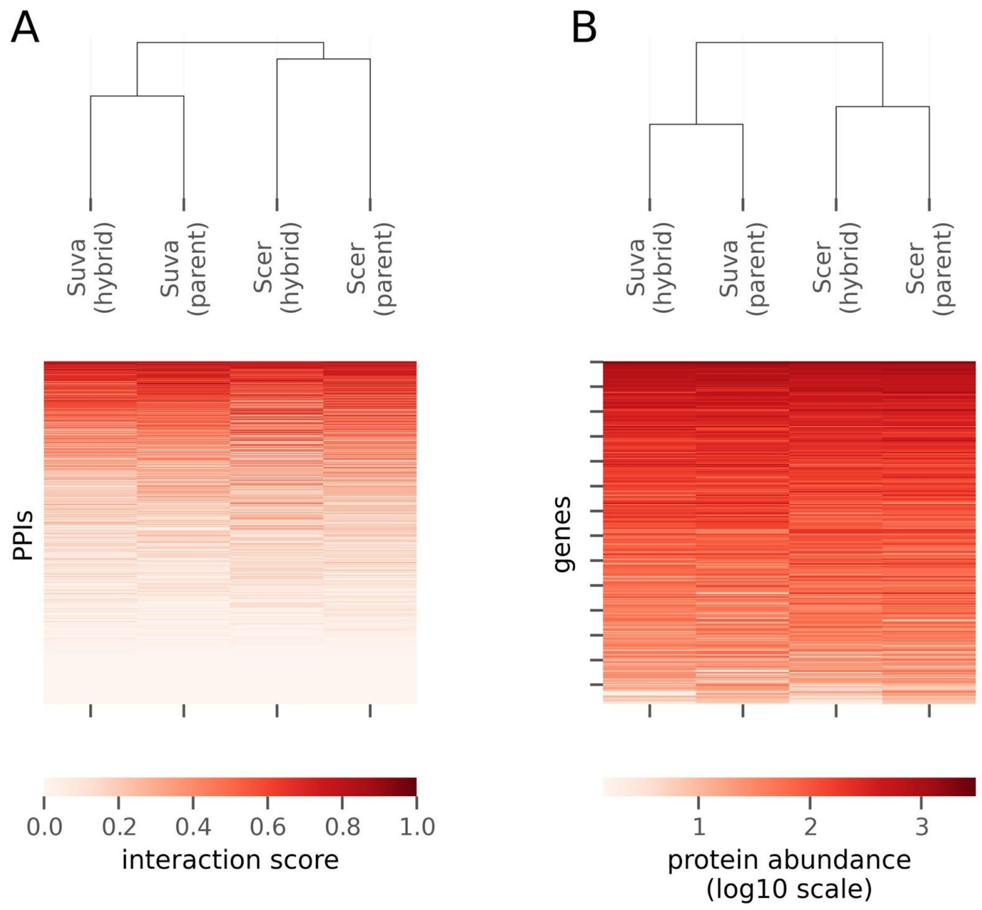


**Figure S6. Clustering based on similarity of interaction scores (A) and protein abundances (B).**

In panel A, interaction scores between pairs of proteins are shown in each row. In panel B, abundance of individual proteins are shown in each row.

The clustering of the interaction types was carried out using Euclidean distances.

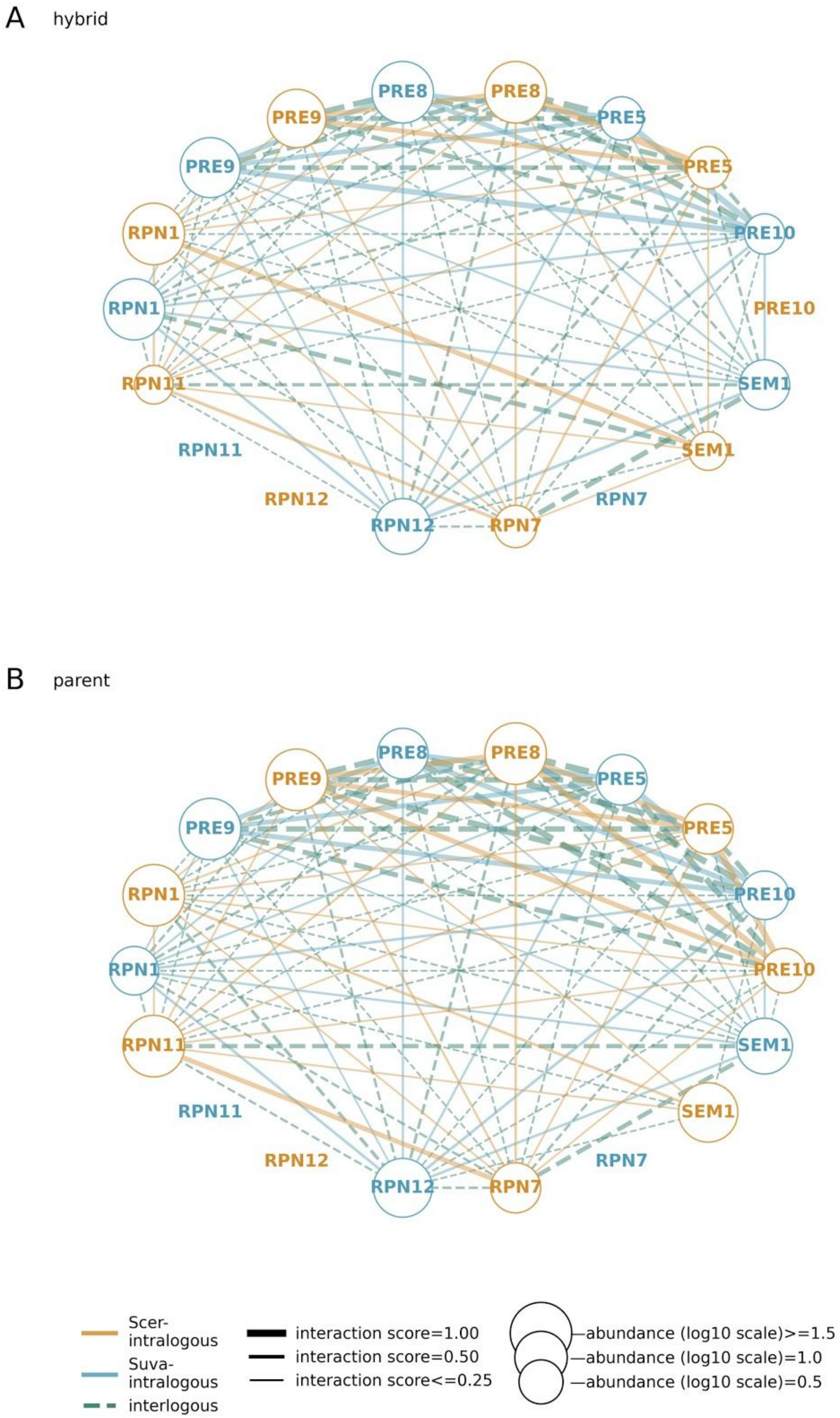
A
hybrid
PRE8
PRE8
PRE9
PRE5
PRE9
PRE5
RPN1
PRE10
RPN1
PRE10
RPN11
SEM1
RPN11
SEM1
RPN12
RPN7
RPN12
RPN7
B
parent
PRE8
PRE8
PRE9
PRE5
PRE9
PRE5
RPN1
PRE10
RPN1
PRE10
RPN11
SEM1
RPN11
SEM1
RPN12
RPN7
RPN12
RPN7
Scer-intralogous
Suva-intralogous
interlogous
interaction score=1.00
interaction score=0.50
interaction score<=0.25
—abundance (log10 scale)>=1.5
—abundance (log10 scale)=1.0
—abundance (log10 scale)=0.5

**Figure S7. PPIs within the 26S proteasome complex.**

**A and B**) PPI networks for the hybrid and parents are shown in panels **A** and **B** respectively. The size of the nodes are in scale with protein abundance and the width of the edges are in scale with interaction scores. Note that the interlogous PPIs in the network of parent species correspond to what PPIs would be in a theoretical hybrid.

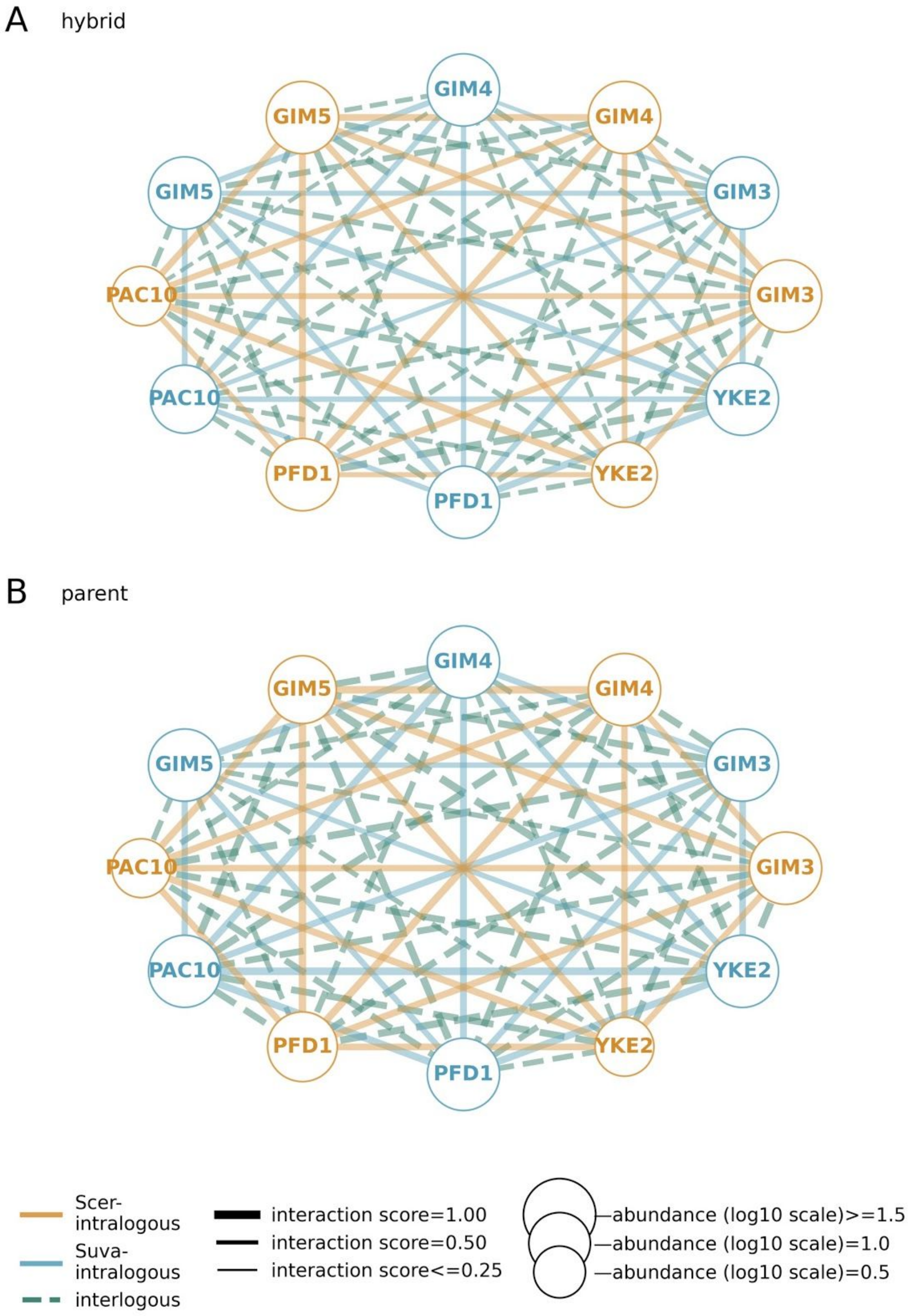


**Figure S8. PPI network for the prefoldin complex, obtained from the comparative proteomics data.**

**A and B**) The PPI networks for the hybrid and parents are shown in panels **A** and **B** respectively. The size of the nodes are in scale with protein abundance and the width of the edges are in scale with interaction scores.Note that the interlogous PPIs in the network of parent species correspond to what PPIs would be in a theoretical hybrid.

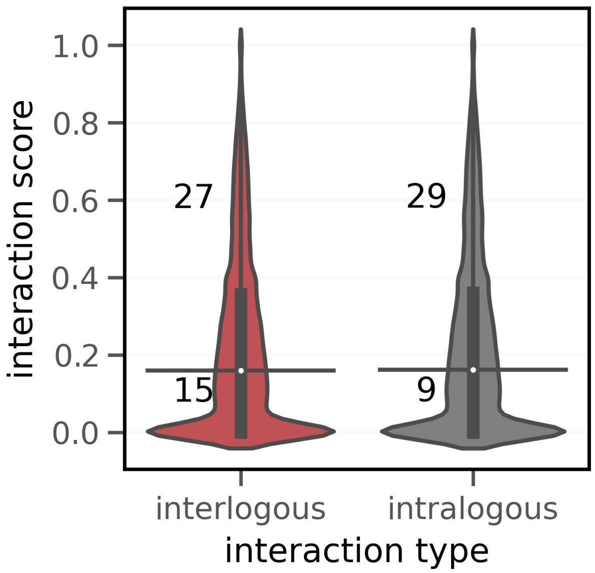


**Figure S9. Comparison of the likely frequencies of chimeric and parental protein complexes in the hybrid.**

Comparison of interaction scores of interlogous and intralogous PPIs in the hybrid. Taking median interaction score as a threshold in each case, the counts of protein complexes that show low and high interaction scores are shown on the violin plots. The medians of the distributions are represented by a horizontal black line and quartiles, by a vertical thick black line.

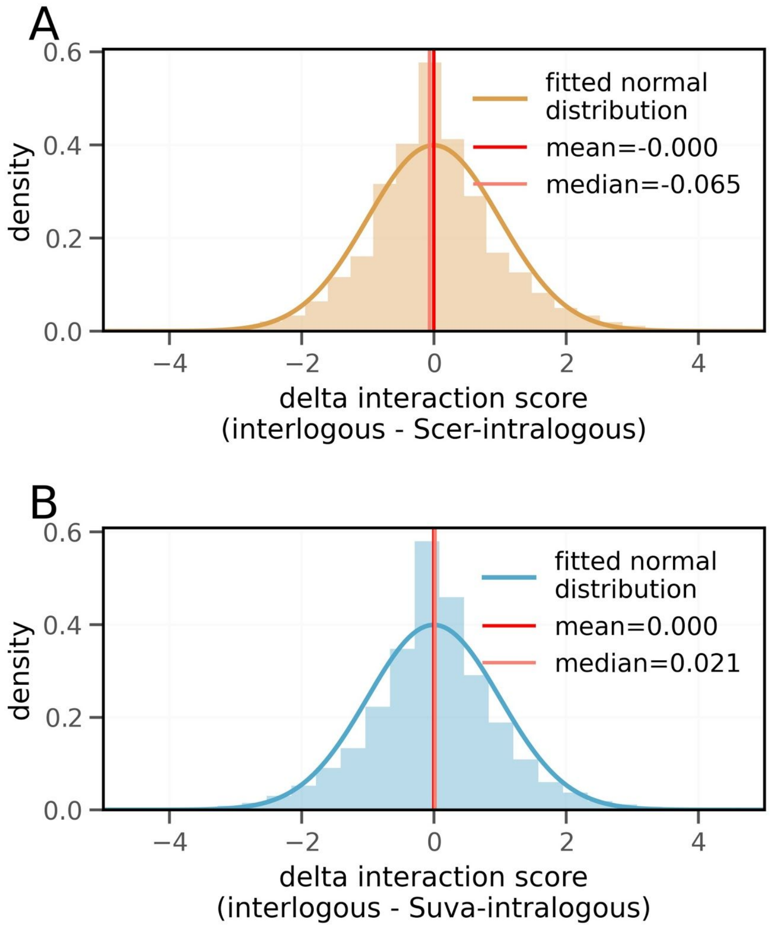


**Figure S10. Analysis of the relative preference for interlogous PPIs over intralogous PPIs in hybrid.**

**A** and **B**) Distributions of the delta interaction score, i.e. ratio between the interlogous and intralogous interaction scores in hybrid, using the z-score scale. The ratio with respect to Scer-intralogous interaction scores is shown in A, while that with respect to Suva-intralogous interaction scores is shown in **B**.

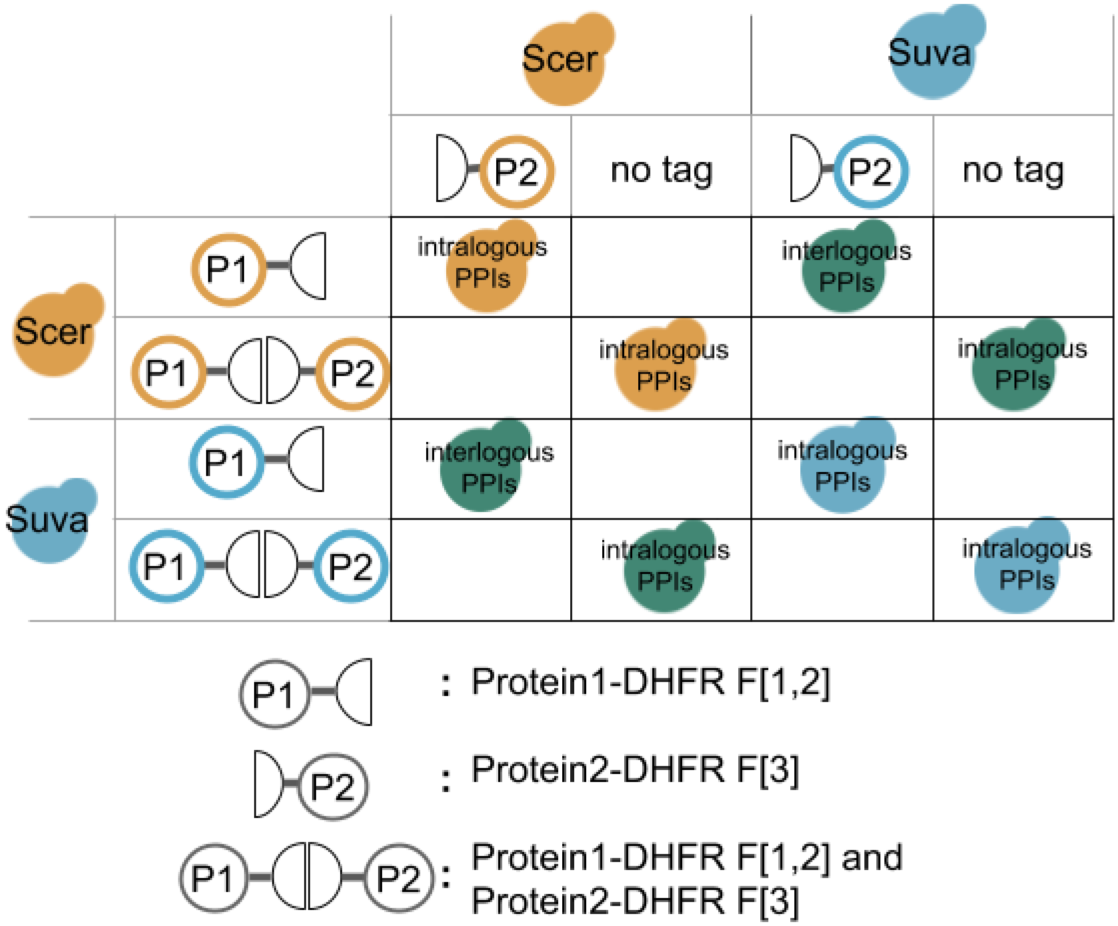


**Figure S11. Strain construction for the DHFR-PCA experiment.**
In order to monitor the interaction between two proteins (P1 and P2), both proteins were tagged with complementary DHFR fragments (DHFR F[1,2] and DHFR F[3]) in haploid cells of opposite mating type. As a control, proteins were tagged with both fragments. Moreover, to allow testing of intralogous PPIs in the hybrid, P1 and P2 were both tagged in the same haploid cells. The crosses with untagged wild-type strains are indicated under the headers labeled as ‘no tag’. PPIs within parents were monitored by crossing between the same species. PPIs within the hybrid were monitored by crossing between the Scer and Suva species. See methods for experimental details.

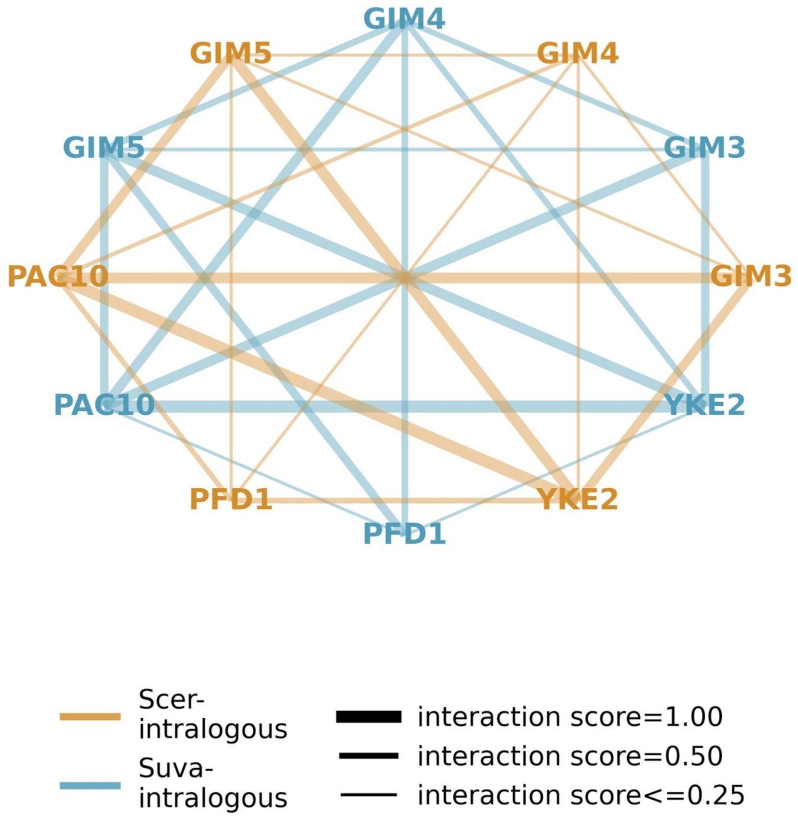


**Figure S12. PPI network within the prefoldin complex in parental yeast species, obtained from DHFR-PCA.**

Intralogous PPIs for Scer interactors are shown in yellow and those for Suva, in blue.

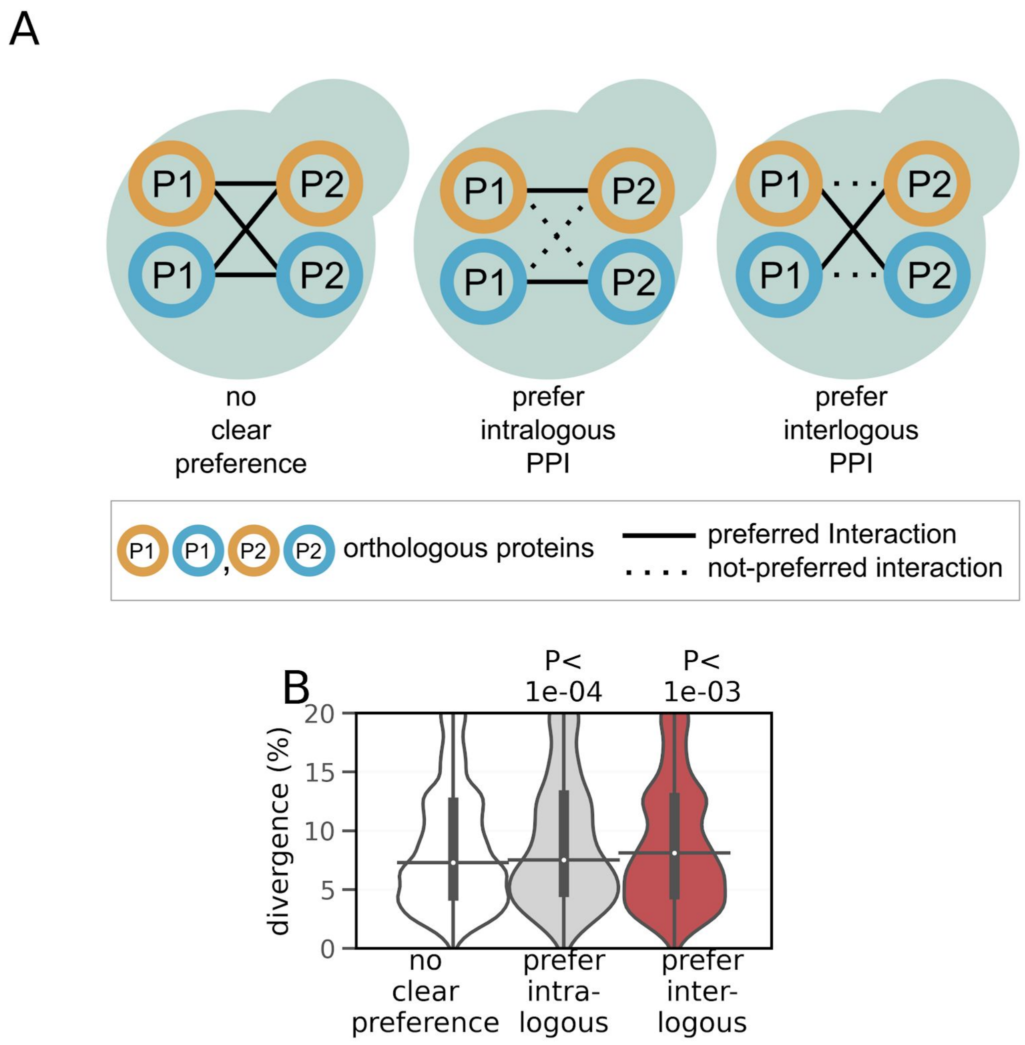


**Figure S13. Preference for intralogous or interlogous PPIs in hybrid is dependent on sequence divergence between orthologous interacting proteins.**

**A**) Schematic representing the hypothesis that preference for either intralogous or interlogous PPIs is dependent on sequence divergence of the orthologous interactors that compete for an interaction.

**B**) Preference for intralogous or interlogous PPIs is associated with higher protein sequence divergence between orthologous interacting proteins (shown on y-axis), compared to PPIs with no preference that are associated with lower divergence. PPIs with delta interaction score (as shown in Figure S10) of less than or equal to -1 were classified as 'prefer intralogous, PPIs with delta interaction score of more than or equal to 1 as 'prefer interlogous, while other PPIs were classified as 'no preference'. P-values from two-sided Mann-Whitney U tests are shown. On the violin plots, the medians of the distributions are represented by a horizontal black line and quartiles, by a vertical thick black line.

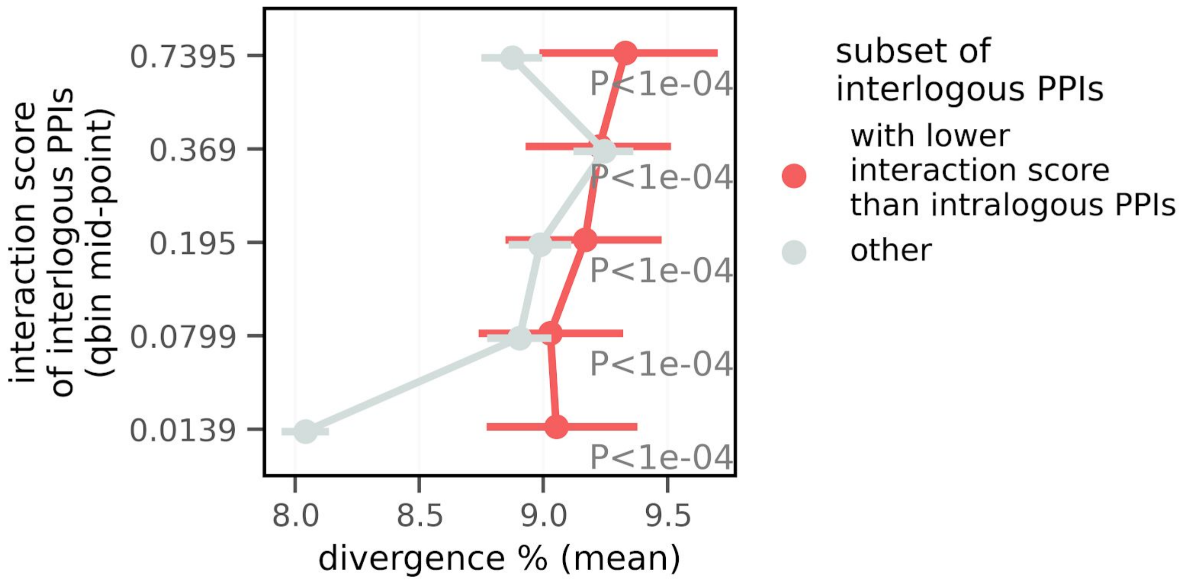


**Figure S14. Divergence of the interlogous PPIs with potentially low compatibility.** Mean divergence of the interactors in interlogous PPIs (x-axis) is compared between two subsets of interlogous PPIs, across the range of interaction scores (y-axis). The subset of interlogous PPIs 'with lower interaction score than intralogous PPIs' in parental species and hybrid represents the interlogous PPIs with the potentially low compatibility. The 'other' subset contains the rest of the interlogous PPIs. For clarity, interaction scores of interlogous PPIs (on y-axis) are divided into 5 bins of equal sizes. Average divergence values per bin is shown (on x-axis) with bars indicating 95 % confidence interval. P-values from two-sided Mann-Whitney U tests are shown.

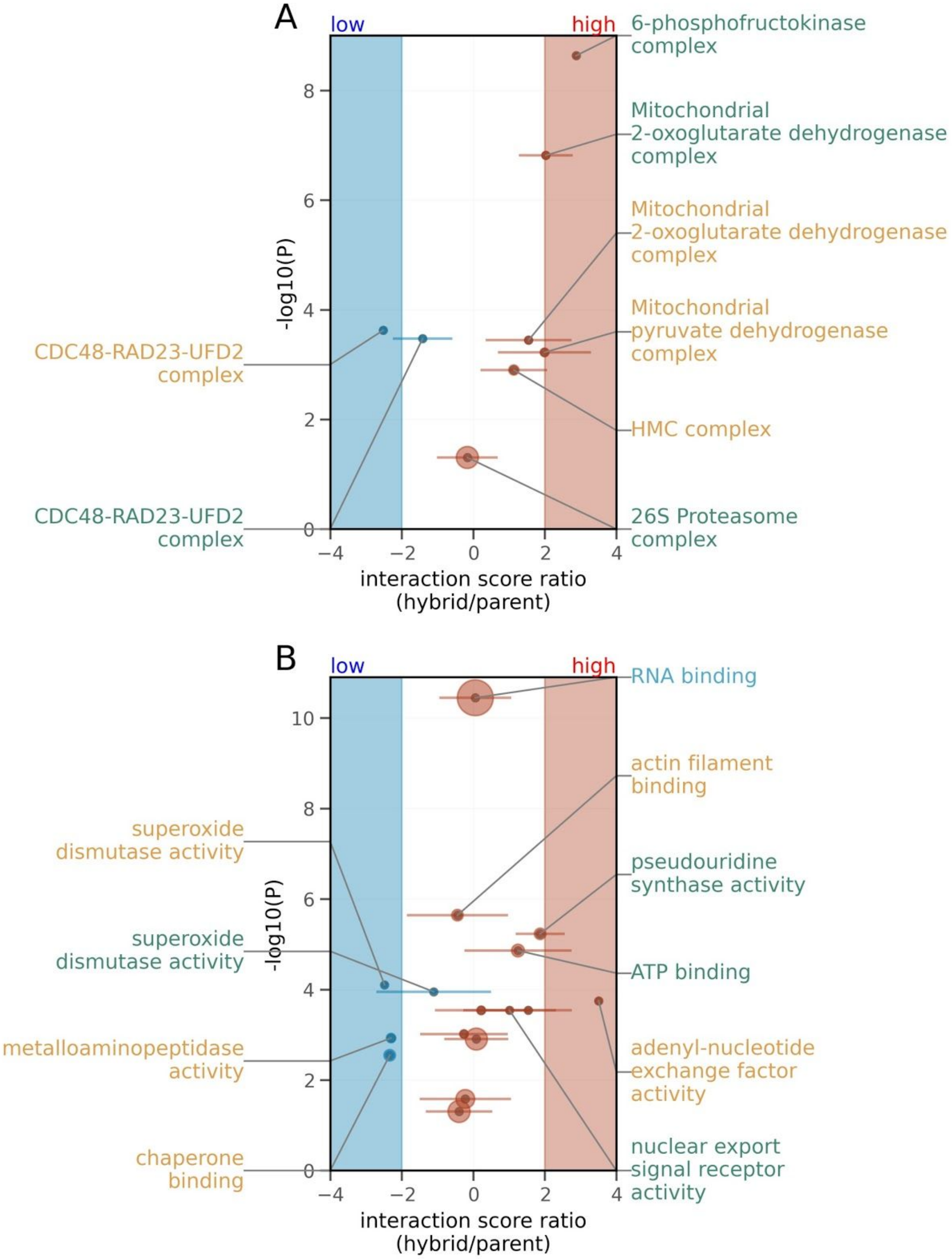


**Figure S15. Gene set enrichment analysis.**

Enrichment of protein complexes (**A**) and GO terms from the Molecular functions (**B**). Volcano plots showing the significantly enriched gene sets.

In both panels, the median value of z-score normalized hybrid/parent ratio of interaction scores, per a set of PPIs corresponding to a gene set is shown on x-axis and -log10(P-value) for enrichment (hypergeometric test, FDR corrected by Benjamini & Hochberg method) is shown on y-axis. Each point corresponds to sets of Scer-intralogous (yellow annotations), Suva-intralogous (blue annotations) or interlogous (green annotations) PPIs. Size of the points is proportional to the number of genes per gene set detected in the proteomics data. For clarity, a maximum of six most significantly low (on left) and significantly high (on right) enrichment gene sets are annotated on the plots. Horizontal error bars indicate standard deviation of the ratio for the set of PPIs. The region shaded in blue indicates a significantly low ratio (z-score ≤ -2), while the region shaded in red indicates a significantly high ratio (z-score ≥ 2).

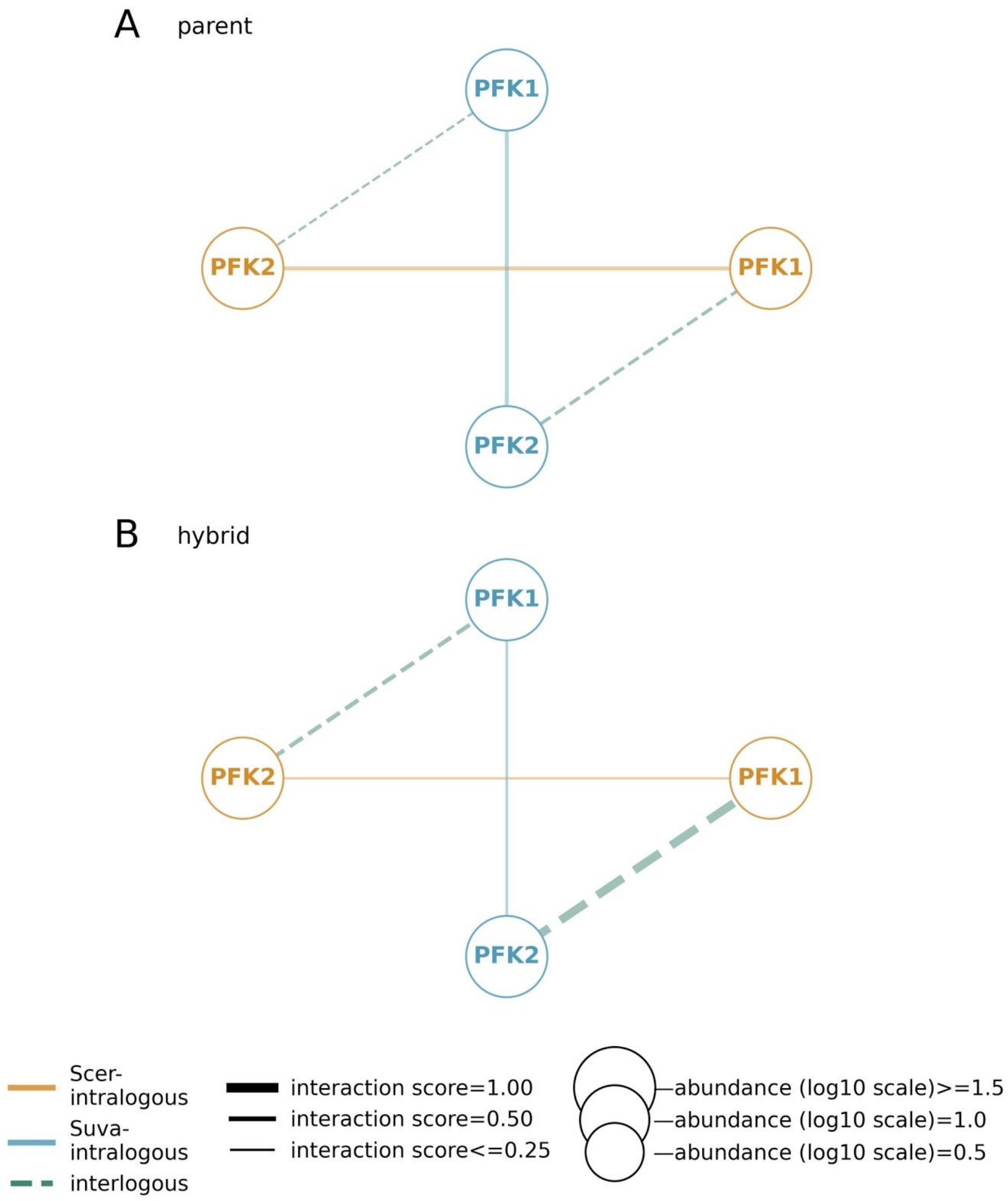


**Figure S16. PPI network of the 6-phosphofructokinase complex.**

The size of the nodes are in scale with protein abundance and the width of the edges are in scale with interaction scores.

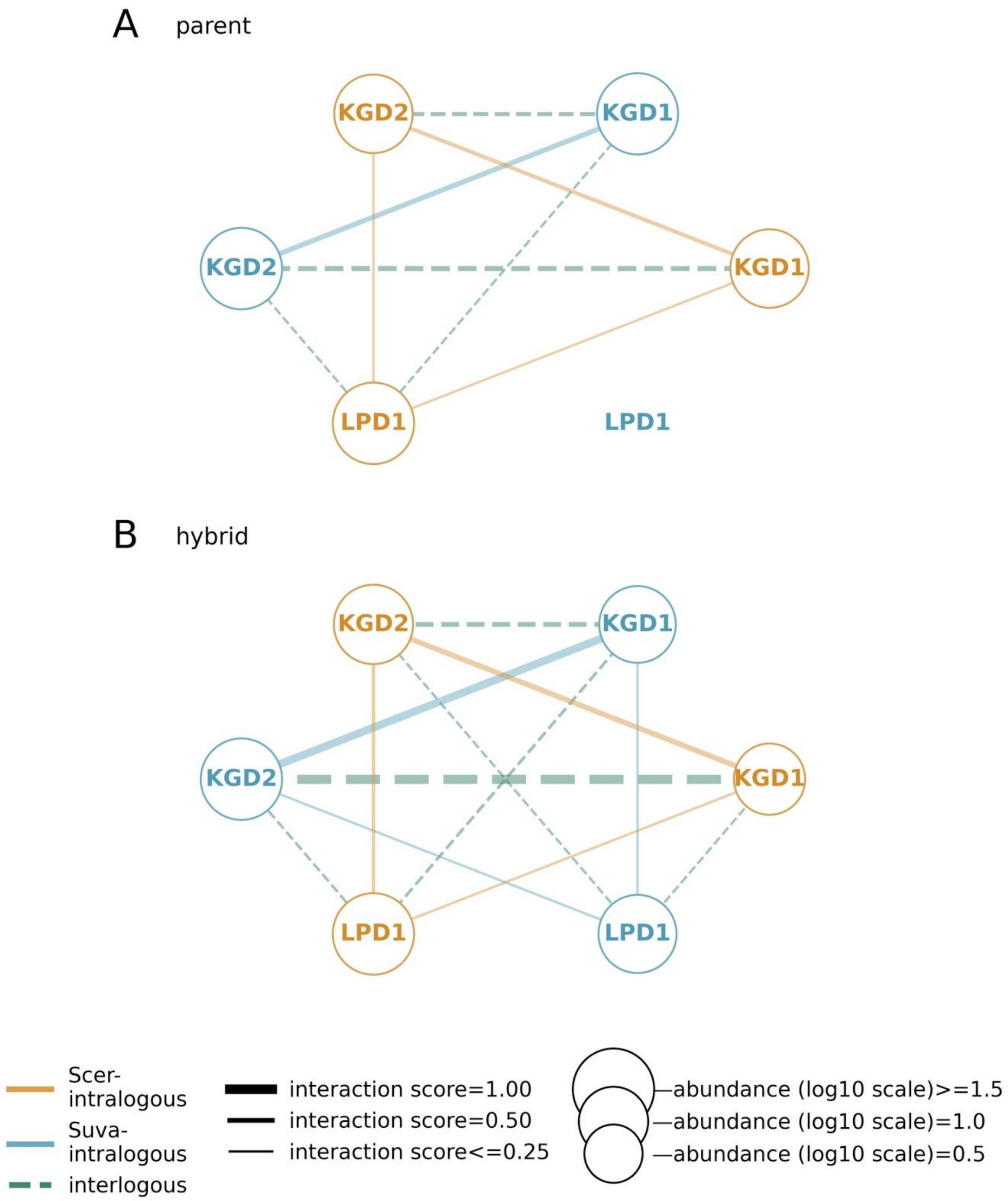


**Figure S17. PPI network of the Mitochondrial 2-oxoglutarate dehydrogenase complex.**

The size of the nodes are in scale with protein abundance and the width of the edges are in scale with interaction scores.

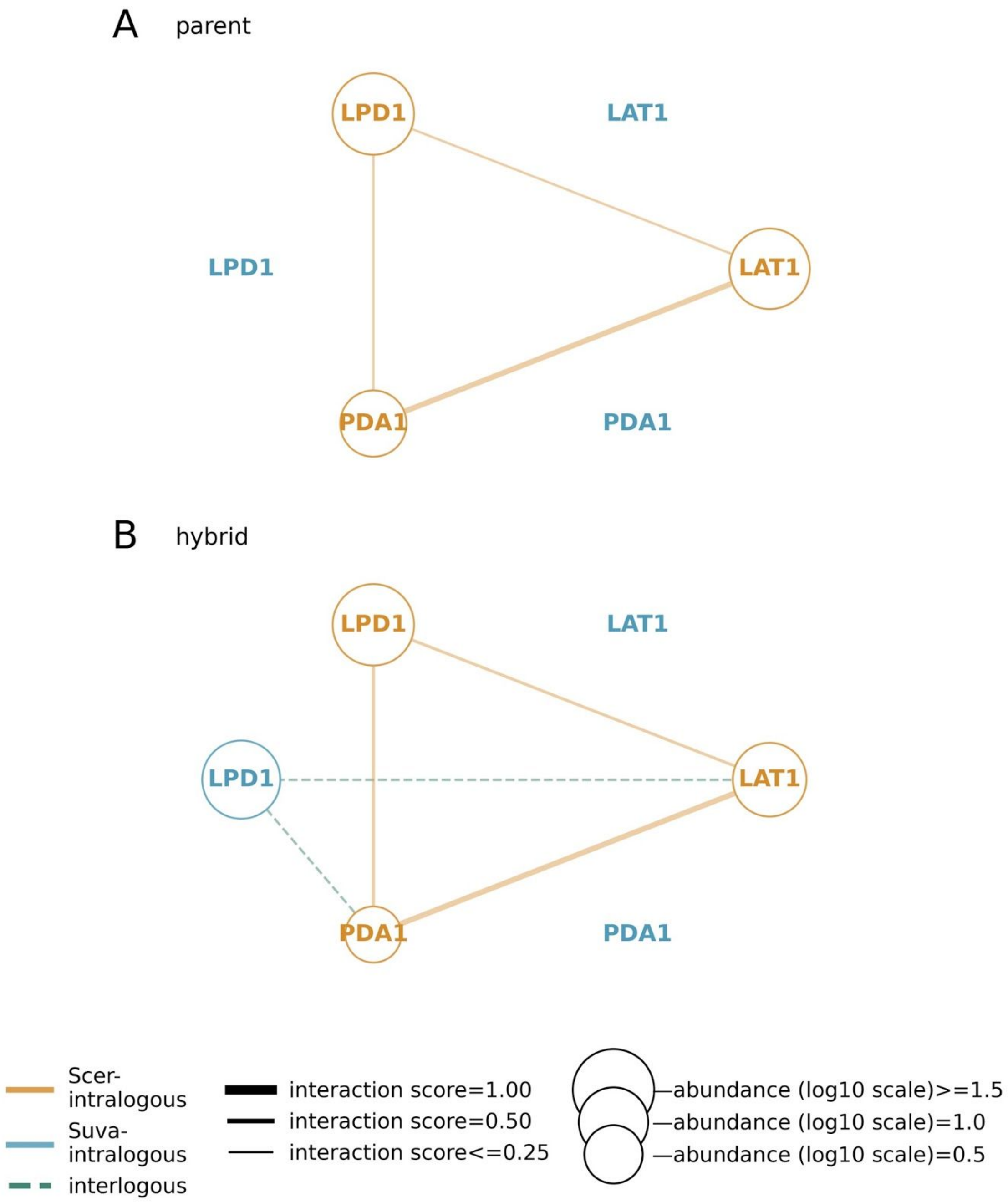


**Figure S18. PPI network of the Mitochondrial pyruvate dehydrogenase complex.** The size of the nodes are in scale with protein abundance and the width of the edges are in scale with interaction scores.The size of the nodes are in the scale of protein abundance, widths of the edges are in the scale of the interaction scores.

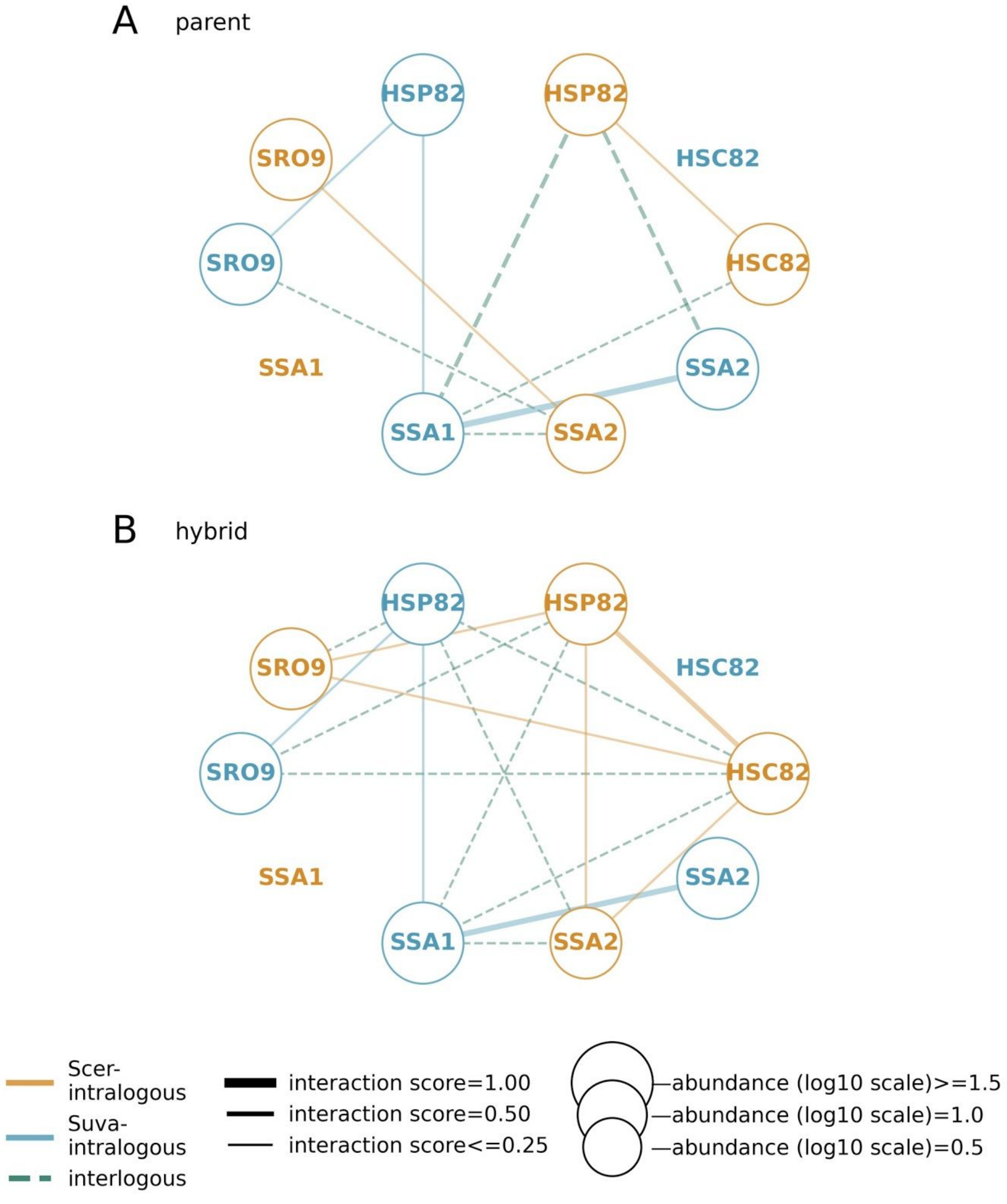


**Figure S19. PPI network of the HMC complex.**

The size of the nodes are in scale with protein abundance and the width of the edges are in scale with interaction scores.The size of the nodes are in the scale of protein abundance, widths of the edges are in the scale of the interaction scores.

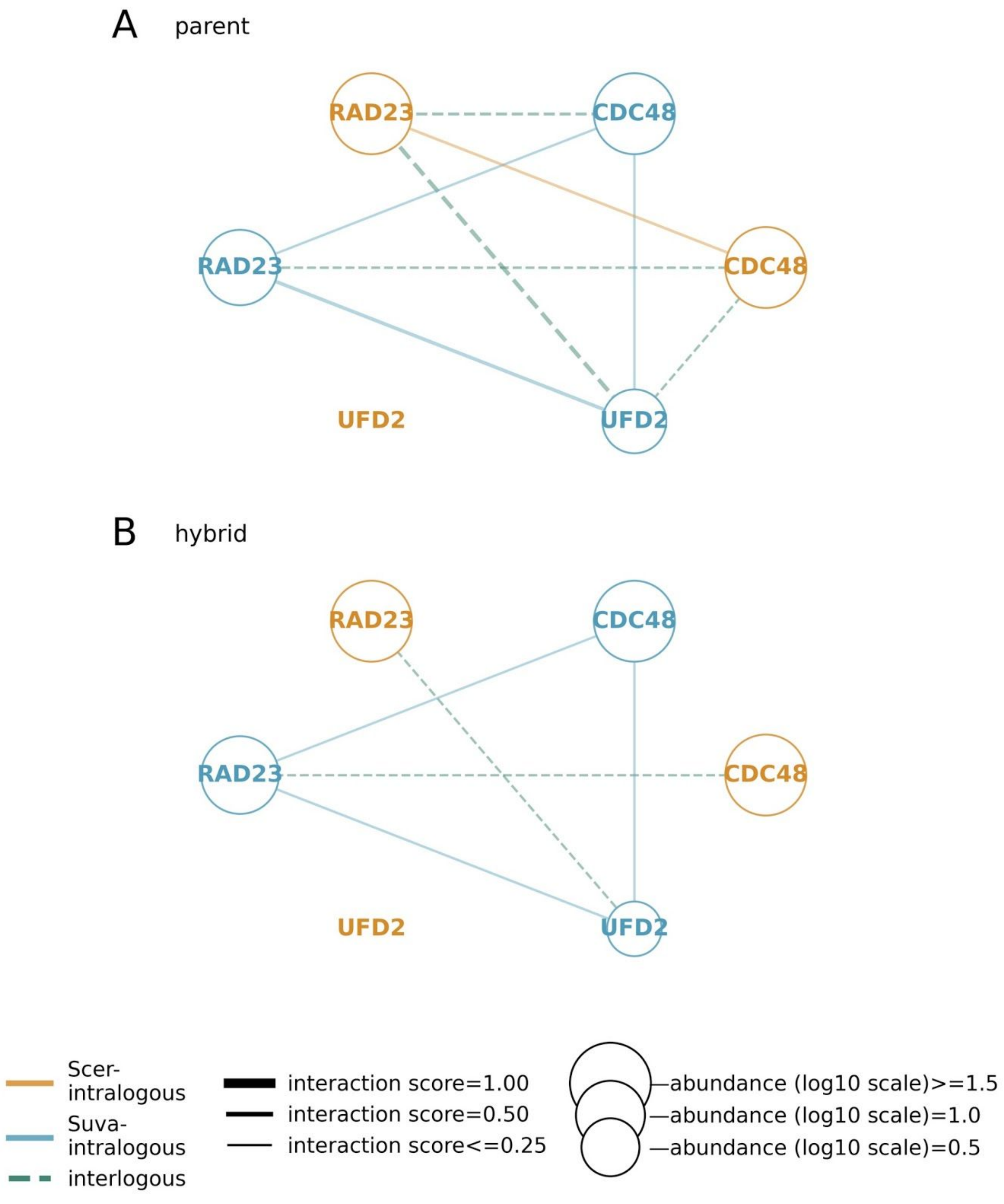


**Figure S20. PPI network of the CDC48-RAD23-UFD2 complex.**

The size of the nodes are in scale with protein abundance and the width of the edges are in scale with interaction scores.The size of the nodes are in the scale of protein abundance, widths of the edges are in the scale of the interaction scores.

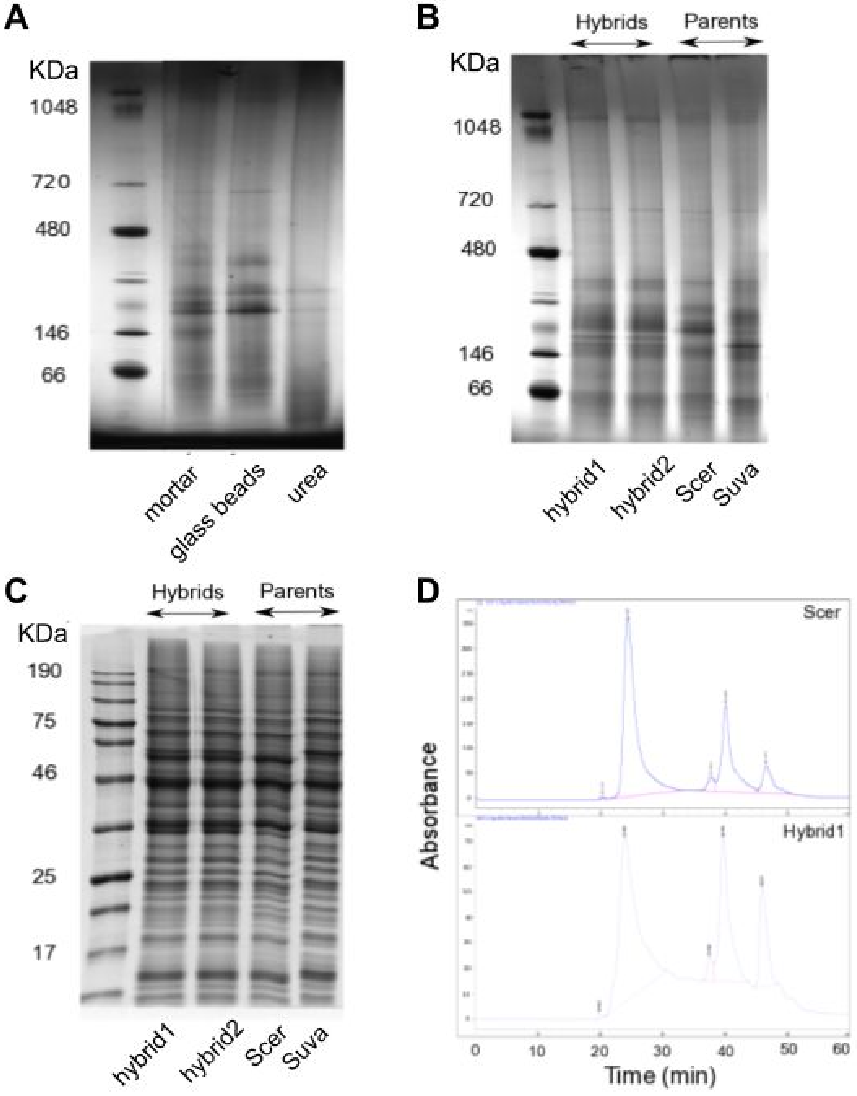


**Figure S21. Experimental optimization of the SEC-PCP-SILAC method in yeast hybrid and parental strains.**

**A)** Native gel which shows comparison of two different lysis protocols. The protocol that shows higher efficiency for protein complex extraction (more bands on gel) was selected (mortar). Urea was used as a control to disrupt protein complexes.

**B)** Native gel which shows comparison of the same protocol (mortar as described in section methods) applied to the different yeast species: Scer, Suva and their hybrids.

**C)** SDS-PAGE gel which shows comparison of the same protocol (mortar as described in section methods) applied to the different yeast species: Scer, Suva and their hybrids.

**D)** Elution profiles obtained after SEC. Profiles obtained for one parent (Scer) and one hybrid are shown only.

### *Supplementary tables*

### Table S1. Description of all the strains that have been used and crossed in this study for the PCP-SILAC and DHFR-PCA experiments.

#### Table S1A. Strains: Mating type and genotype of all haploid strains used in this study including the experiment(s) they were used for.

| **Name** | **Original Strain** | **Species** | **Mating type** | **Genotype** | **Reference** | **Experiment (PCP-SILAC, DHFR-PCA or both)** |
|---|---|---|---|---|---|---|
| BY4741 | S288C | *Saccharomyces cerevisiae* | MAT**a** | *his3 leu2 ura3 met15* | Brachmann, C.B., et al., Designer deletion strains derived from *Saccharomyces cerevisiae* S288C: A useful set of strains and plasmids for PCR-mediated gene disruption and other applications. Yeast, 1998. 14(2): p. 115-132. | DHFR-PCA |
| BY4741-*lys2Δ* | S288C | *Saccharomyces cerevisiae* | MAT**a** | *lys2::KANMX4 his3 leu2 ura3 met15* | Yeast Deletion Collection | PCP-SILAC |
| BY4742 | S288C | *Saccharomyces cerevisiae* | MAT**α** | *his3 leu2 ura3 lys2* | Brachmann, C.B., et al., Designer deletion strains derived from *Saccharomyces cerevisiae* S288C: A useful set of strains and plasmids for PCR-mediated gene disruption and other applications. Yeast, 1998. 14(2): p. 115-132. | Both |
| MG032 | CBS7001 | *Saccharomyces uvarum* | MAT**a** | *ho::URA3 his3 ura3 trp lys2* | Leducq, J.-B., et al., Evidence for the Robustness of Protein Complexes to interspecies Hybridization (Protein Complexes in Hybrids). Protein Complexes in Hybrids, 2012. 8(12): p. e1003161. | PCP-SILAC |
| MG031 | CBS7001 | *Saccharomyces uvarum* | MAT**α** | *ho::URA3 his3 ura3 ade2 lys2* | Leducq, J.-B., et al., Evidence for the Robustness of Protein Complexes to interspecies Hybridization (Protein Complexes in Hybrids). Protein Complexes in Hybrids, 2012. 8(12): p. e1003161. | DHFR-PCA |
| JBL033 | CBS7001 | *Saccharomyces uvarum* | MAT**α** | *hoΔ his3 ura3 trp lys2* | Leducq, J.-B., et al., Evidence for the Robustness of Protein Complexes to interspecies Hybridization (Protein Complexes in Hybrids). Protein Complexes in Hybrids, 2012. 8(12): p. e1003161. | PCP-SILAC |
| GIM3 F[1,2] | BY4741 (S288C) | *Saccharomyces cerevisiae* | MAT**a** | *GIM3-DHFR F[1,2]-NATMX6 hoΔ his3 leu2 ura3 met15* | Yeast Protein Interactome Collection | DHFR-PCA |
| GIM4 F[1,2] | BY4741 (S288C) | *Saccharomyces cerevisiae* | MAT**a** | *GIM4-DHFR F[1,2]-NATMX6 hoΔ his3 leu2 ura3 met15* | Yeast Protein Interactome Collection | DHFR-PCA |
| GIM5 F[1,2] | BY4741 (S288C) | *Saccharomyces cerevisiae* | MAT**a** | *GIM5-DHFR F[1,2]-NATMX6 hoΔ his3 leu2 ura3 met15* | Yeast Protein Interactome Collection | DHFR-PCA |
| PAC10 F[1,2] | BY4741 (S288C) | *Saccharomyces cerevisiae* | MAT**a** | *PAC10-DHFR F[1,2]-NATMX6 hoΔ his3 leu2 ura3 met15* | Yeast Protein Interactome Collection | DHFR-PCA |
| PFD1 F[1,2] | BY4741 (S288C) | *Saccharomyces cerevisiae* | MAT**a** | *PFD1-DHFR F[1,2]-NATMX6 hoΔ his3 leu2 ura3 met15* | Yeast Protein Interactome Collection | DHFR-PCA |
| YKE2 F[1,2] | BY4741 (S288C) | *Saccharomyces cerevisiae* | MAT**a** | *YKE2-DHFR F[1,2]-NATMX6 hoΔ his3 leu2 ura3 met15* | Yeast Protein Interactome Collection | DHFR-PCA |
| GIM3 F[3] | BY4742 (S288C) | *Saccharomyces cerevisiae* | MAT**α** | *GIM3-DHFR F[3]-HPHMX6 hoΔ his3 leu2 ura3 lys2* | Yeast Protein Interactome Collection | DHFR-PCA |
| GIM4 F[3] | BY4742 (S288C) | *Saccharomyces cerevisiae* | MAT**α** | *GIM4-DHFR F[3]-HPHMX6 hoΔ his3 leu2 ura3 lys2* | Yeast Protein Interactome Collection | DHFR-PCA |
| GIM5 F[3] | BY4742 (S288C) | *Saccharomyces cerevisiae* | MAT**α** | *GIM5-DHFR F[3]-HPHMX6 hoΔ his3 leu2 ura3 lys2* | Yeast Protein Interactome Collection | DHFR-PCA |
| PAC10 F[3] | BY4742 (S288C) | *Saccharomyces cerevisiae* | MAT**α** | *PAC10-DHFR F[3]-HPHMX6 hoΔ his3 leu2 ura3 lys2* | Yeast Protein Interactome Collection | DHFR-PCA |

| | | | | | | |
|---|---|---|---|---|---|---|
| CAB029 | BY4742 (S288C) | *Saccharomyces cerevisiae* | MAT**α** | *PFD1-DHFR F[3]-HPHMX6 hoΔ his3 leu2 ura3 lys2* | This study | DHFR-PCA |
| YKE2 F[3] | BY4742 (S288C) | *Saccharomyces cerevisiae* | MAT**α** | *YKE2-DHFR F[3]-HPHMX6 hoΔ his3 leu2 ura3 lys2* | Yeast Protein Interactome Collection | DHFR-PCA |
| CAB020 | MG032 (CBS7001) | *Saccharomyces uvarum* | MAT**a** | *GIM3-DHFR F[1,2]-NATMX6 ho::URA3 his3 ura3 trp lys2* | This study | DHFR-PCA |
| CAB021 | MG032 (CBS7001) | *Saccharomyces uvarum* | MAT**a** | *GIM4-DHFR F[1,2]-NATMX6 ho::URA3 his3 ura3 trp lys2* | This study | DHFR-PCA |
| CAB022 | MG032 (CBS7001) | *Saccharomyces uvarum* | MAT**a** | *GIM5-DHFR F[1,2]-NATMX6 ho::URA3 his3 ura3 trp lys2* | This study | DHFR-PCA |
| CAB023 | MG032 (CBS7001) | *Saccharomyces uvarum* | MAT**a** | *PAC10-DHFR F[1,2]-NATMX6 ho::URA3 his3 ura3 trp lys2* | This study | DHFR-PCA |
| CAB046 | MG032 (CBS7001) | *Saccharomyces uvarum* | MAT**a** | *PFD1-DHFR F[1,2]-NATMX6 ho::URA3 his3 ura3 trp lys2* | This study | DHFR-PCA |
| CAB024 | MG032 (CBS7001) | *Saccharomyces uvarum* | MAT**a** | *YKE2-DHFR F[1,2]-NATMX6 ho::URA3 his3 ura3 trp lys2* | This study | DHFR-PCA |
| CAB039 | MG031 (CBS7001) | *Saccharomyces uvarum* | MAT**α** | *GIM3-DHFR F[3]-HPHMX6 ho::URA3 his3 ura3 ade2 lys2* | This study | DHFR-PCA |
| CAB040 | MG031 (CBS7001) | *Saccharomyces uvarum* | MAT**α** | *GIM4-DHFR F[3]-HPHMX6 ho::URA3 his3 ura3 ade2 lys2* | This study | DHFR-PCA |
| CAB041 | MG031 (CBS7001) | *Saccharomyces uvarum* | MAT**α** | *GIM5-DHFR F[3]-HPHMX6 ho::URA3 his3 ura3 ade2 lys2* | This study | DHFR-PCA |
| CAB042 | MG031 (CBS7001) | *Saccharomyces uvarum* | MAT**α** | *PAC10-DHFR F[3]-HPHMX6 ho::URA3 his3 ura3 ade2 lys2* | This study | DHFR-PCA |
| CAB038 | MG031 (CBS7001) | *Saccharomyces uvarum* | MAT**α** | *PFD1-DHFR F[3]-HPHMX6 ho::URA3 his3 ura3 ade2 lys2* | This study | DHFR-PCA |
| CAB043 | MG031 (CBS7001) | *Saccharomyces uvarum* | MAT**α** | *YKE2-DHFR F[3]-HPHMX6 ho::URA3 his3 ura3 ade2 lys2* | This study | DHFR-PCA |
| CAB047 | BY4741 (S288C) | *Saccharomyces cerevisiae* | MAT**a** | *PFD1-DHFR F[1,2]-NATMX6 GIM3-DHFR F[3]-HPHMX6 hoΔ his3 leu2 ura3 met15* | This study | DHFR-PCA |
| CAB048 | BY4741 (S288C) | *Saccharomyces cerevisiae* | MAT**a** | *PFD1-DHFR F[1,2]-NATMX6 GIM4-DHFR F[3]-HPHMX6 hoΔ his3 leu2 ura3 met15* | This study | DHFR-PCA |
| CAB049 | BY4741 (S288C) | *Saccharomyces cerevisiae* | MAT**a** | *PFD1-DHFR F[1,2]-NATMX6 GIM5-DHFR F[3]-HPHMX6 hoΔ his3 leu2 ura3 met15* | This study | DHFR-PCA |
| CAB050 | BY4741 (S288C) | *Saccharomyces cerevisiae* | MAT**a** | *PFD1-DHFR F[1,2]-NATMX6 PAC10-DHFR F[3]-HPHMX6 hoΔ his3 leu2 ura3 met15* | This study | DHFR-PCA |
| CAB051 | BY4741 (S288C) | *Saccharomyces cerevisiae* | MAT**a** | *PFD1-DHFR F[1,2]-NATMX6 YKE2-DHFR F[3]-HPHMX6 hoΔ his3 leu2 ura3 met15* | This study | DHFR-PCA |
| CAB052 | BY4741 (S288C) | *Saccharomyces cerevisiae* | MAT**a** | *GIM3-DHFR F[1,2]-NATMX6* | This study | DHFR-PCA |

| | | | | | | |
|---|---|---|---|---|---|---|
| | | | | *GIM4-DHFR F[3]-HPHMX6 hoΔ his3 leu2 ura3 met15* | | |
| CAB053 | BY4741 (S288C) | *Saccharomyces cerevisiae* | MAT**a** | *GIM3-DHFR F[1,2]-NATMX6 GIM5-DHFR F[3]-HPHMX6 hoΔ his3 leu2 ura3 met15* | This study | DHFR-PCA |
| CAB054 | BY4741 (S288C) | *Saccharomyces cerevisiae* | MAT**a** | *GIM3-DHFR F[1,2]-NATMX6 PAC10-DHFR F[3]-HPHMX6 hoΔ his3 leu2 ura3 met15* | This study | DHFR-PCA |
| CAB055 | BY4741 (S288C) | *Saccharomyces cerevisiae* | MAT**a** | *GIM3-DHFR F[1,2]-NATMX6 YKE2-DHFR F[3]-HPHMX6 hoΔ his3 leu2 ura3 met15* | This study | DHFR-PCA |
| CAB056 | BY4741 (S288C) | *Saccharomyces cerevisiae* | MAT**a** | *GIM4-DHFR F[1,2]-NATMX6 GIM5-DHFR F[3]-HPHMX6 hoΔ his3 leu2 ura3 met15* | This study | DHFR-PCA |
| CAB057 | BY4741 (S288C) | *Saccharomyces cerevisiae* | MAT**a** | *GIM4-DHFR F[1,2]-NATMX6 PAC10-DHFR F[3]-HPHMX6 hoΔ his3 leu2 ura3 met15* | This study | DHFR-PCA |
| CAB058 | BY4741 (S288C) | *Saccharomyces cerevisiae* | MAT**a** | *GIM4-DHFR F[1,2]-NATMX6 YKE2-DHFR F[3]-HPHMX6 hoΔ his3 leu2 ura3 met15* | This study | DHFR-PCA |
| CAB059 | BY4741 (S288C) | *Saccharomyces cerevisiae* | MAT**a** | *GIM5-DHFR F[1,2]-NATMX6 PAC10-DHFR F[3]-HPHMX6 hoΔ his3 leu2 ura3 met15* | This study | DHFR-PCA |
| CAB060 | BY4741 (S288C) | *Saccharomyces cerevisiae* | MAT**a** | *GIM5-DHFR F[1,2]-NATMX6 YKE2-DHFR F[3]-HPHMX6 hoΔ his3 leu2 ura3 met15* | This study | DHFR-PCA |
| CAB061 | BY4741 (S288C) | *Saccharomyces cerevisiae* | MAT**a** | *PAC10-DHFR F[1,2]-NATMX6 YKE2-DHFR F[3]-HPHMX6 hoΔ his3 leu2 ura3 met15* | This study | DHFR-PCA |
| CAB062 | MG032 (CBS7001) | *Saccharomyces uvarum* | MAT**a** | *PFD1-DHFR F[1,2]-NATMX6 GIM3-DHFR F[3]-HPHMX6 ho::URA3 his3 ura3 trp lys2* | This study | DHFR-PCA |
| CAB063 | MG032 (CBS7001) | *Saccharomyces uvarum* | MAT**a** | *PFD1-DHFR F[1,2]-NATMX6 GIM4-DHFR F[3]-HPHMX6 ho::URA3 his3 ura3 trp lys2* | This study | DHFR-PCA |
| CAB064 | MG032 (CBS7001) | *Saccharomyces uvarum* | MAT**a** | *PFD1-DHFR F[1,2]-NATMX6 GIM5-DHFR F[3]-HPHMX6 ho::URA3 his3 ura3 trp lys2* | This study | DHFR-PCA |
| CAB065 | MG032 (CBS7001) | *Saccharomyces uvarum* | MAT**a** | *PFD1-DHFR F[1,2]-NATMX6 PAC10-DHFR F[3]-HPHMX6 ho::URA3 his3 ura3 trp lys2* | This study | DHFR-PCA |
| CAB066 | MG032 (CBS7001) | *Saccharomyces uvarum* | MAT**a** | *PFD1-DHFR F[1,2]-NATMX6 YKE2-DHFR F[3]-HPHMX6 ho::URA3 his3 ura3 trp lys2* | This study | DHFR-PCA |
| CAB067 | MG032 (CBS7001) | *Saccharomyces uvarum* | MAT**a** | *GIM3-DHFR F[1,2]-NATMX6 GIM4-DHFR* | This study | DHFR-PCA |

| | | | | | | |
|---|---|---|---|---|---|---|
| | | | | *F[3]-HPHMX6 ho::URA3 his3 ura3 trp lys2* | | |
| CAB068 | MG032 (CBS7001) | *Saccharomyces uvarum* | MAT**a** | *GIM3-DHFR F[1,2]-NATMX6 GIM5-DHFR F[3]-HPHMX6 ho::URA3 his3 ura3 trp lys2* | This study | DHFR-PCA |
| CAB069 | MG032 (CBS7001) | *Saccharomyces uvarum* | MAT**a** | *GIM3-DHFR F[1,2]-NATMX6 PAC10-DHFR F[3]-HPHMX6 ho::URA3 his3 ura3 trp lys2* | This study | DHFR-PCA |
| CAB070 | MG032 (CBS7001) | *Saccharomyces uvarum* | MAT**a** | *GIM3-DHFR F[1,2]-NATMX6 YKE2-DHFR F[3]-HPHMX6 ho::URA3 his3 ura3 trp lys2* | This study | DHFR-PCA |
| CAB071 | MG032 (CBS7001) | *Saccharomyces uvarum* | MAT**a** | *GIM4-DHFR F[1,2]-NATMX6 GIM5-DHFR F[3]-HPHMX6 ho::URA3 his3 ura3 trp lys2* | This study | DHFR-PCA |
| CAB072 | MG032 (CBS7001) | *Saccharomyces uvarum* | MAT**a** | *GIM4-DHFR F[1,2]-NATMX6 PAC10-DHFR F[3]-HPHMX6 ho::URA3 his3 ura3 trp lys2* | This study | DHFR-PCA |
| CAB073 | MG032 (CBS7001) | *Saccharomyces uvarum* | MAT**a** | *GIM4-DHFR F[1,2]-NATMX6 YKE2-DHFR F[3]-HPHMX6 ho::URA3 his3 ura3 trp lys2* | This study | DHFR-PCA |
| CAB074 | MG032 (CBS7001) | *Saccharomyces uvarum* | MAT**a** | *GIM5-DHFR F[1,2]-NATMX6 PAC10-DHFR F[3]-HPHMX6 ho::URA3 his3 ura3 trp lys2* | This study | DHFR-PCA |
| CAB075 | MG032 (CBS7001) | *Saccharomyces uvarum* | MAT**a** | *GIM5-DHFR F[1,2]-NATMX6 YKE2-DHFR F[3]-HPHMX6 ho::URA3 his3 ura3 trp lys2* | This study | DHFR-PCA |
| CAB076 | MG032 (CBS7001) | *Saccharomyces uvarum* | MAT**a** | *PAC10-DHFR F[1,2]-NATMX6 YKE2-DHFR F[3]-HPHMX6 ho::URA3 his3 ura3 trp lys2* | This study | DHFR-PCA |
| LSM8 F[1,2] | BY4741 (S288C) | *Saccharomyces cerevisiae* | MAT**a** | *LSM8-DHFR F[1,2]-NATMX6 hoΔ his3 leu2 ura3 met15* | Yeast Protein Interactome Collection | DHFR-PCA |
| CDC39 F[3] | BY4742 (S288C) | *Saccharomyces cerevisiae* | MAT**α** | *CDC39-DHFR F[3]-HPHMX6 hoΔ his3 leu2 ura3 lys2* | Yeast Protein Interactome Collection | DHFR-PCA |

### Table S1B. PCP-SILAC: Crosses made for the PCP-SILAC experiment.

| Cross | MATa strain | MATα strain |
|---|---|---|
| Scer | BY4741-*lys2Δ* | BY4742 |
| Suva | MG032 | JBL033 |
| hybrid (H1) | BY4741-*lys2Δ* | JBL033 |
| hydrid (H2) | MG032 | BY4742 |

**Table S1C. SILAC labelling combinations used during the SEC-PCP-SILAC experiment.**

| Experiment id | Comparison | L | M | H | Number of technical replicates |
|---|---|---|---|---|---|
| A | parent Scer vs H1 | parent Scer + parent Suva | parent Scer | hybrid 1 H1 | 2 |
| Reciprocal replicate of A | parent Scer vs H2 | parent Scer + parent Suva | parent Scer | hybrid 2 H2 | 2 |
| B | parent Suva vs H1 | parent Scer + parent Suva | parent Suva | hybrid 1 H1 | 2 |
| Reciprocal replicate of B | parent Suva vs H2 | parent Scer + parent Suva | parent Suva | hybrid 2 H2 | 2 |
| C | parent Scer vs parent Suva | parent Scer + parent Suva | parent Scer | parent Suva | 1 |
| Biological replicate of C | parent Scer vs parent Suva | parent Scer + parent Suva | parent Scer | parent Suva | 1 |
| | | | | | **Total 10** |

Notes

- Technical replicates started from the same cell cultures but samples were divided in two before injection for SEC.
- Reciprocal biological replicates started from different independent cultures

**Table S1D. DHFR-PCA: Description of all the crosses made for the DHFR-PCA experiment and identification of the medium used for diploid selection.**

| Cross | PPI type | MATa strain | MATα strain | Gene-DHFR F[1,2] | Gene-DHFR F[3] | Selective medium |
|---|---|---|---|---|---|---|
| Scer | Positive control | LSM8 F[1,2] | CDC39 F[3] | LSM8 | CDC39 | SC -met -lys |
| Scer | Scer-intralogous | GIM3 F[1,2] | GIM4 F[3] | GIM3 | GIM4 | SC -met -lys |
| Scer | Scer-intralogous | GIM3 F[1,2] | GIM5 F[3] | GIM3 | GIM5 | SC -met -lys |
| Scer | Scer-intralogous | GIM3 F[1,2] | PAC10 F[3] | GIM3 | PAC10 | SC -met -lys |
| Scer | Scer-intralogous | GIM3 F[1,2] | CAB029 | GIM3 | PFD1 | SC -met -lys |
| Scer | Scer-intralogous | GIM3 F[1,2] | YKE2 F[3] | GIM3 | YKE2 | SC -met -lys |
| Scer | Scer-intralogous | GIM4 F[1,2] | GIM3 F[3] | GIM4 | GIM3 | SC -met -lys |
| Scer | Scer-intralogous | GIM4 F[1,2] | GIM5 F[3] | GIM4 | GIM5 | SC -met -lys |
| Scer | Scer-intralogous | GIM4 F[1,2] | PAC10 F[3] | GIM4 | PAC10 | SC -met -lys |
| Scer | Scer-intralogous | GIM4 F[1,2] | CAB029 | GIM4 | PFD1 | SC -met -lys |
| Scer | Scer-intralogous | GIM4 F[1,2] | YKE2 F[3] | GIM4 | YKE2 | SC -met -lys |

| | | | | | | |
|---|---|---|---|---|---|---|
| Scer | Scer-intralogous | GIM5 F[1,2] | GIM3 F[3] | GIM5 | GIM3 | SC -met -lys |
| Scer | Scer-intralogous | GIM5 F[1,2] | GIM4 F[3] | GIM5 | GIM4 | SC -met -lys |
| Scer | Scer-intralogous | GIM5 F[1,2] | PAC10 F[3] | GIM5 | PAC10 | SC -met -lys |
| Scer | Scer-intralogous | GIM5 F[1,2] | CAB029 | GIM5 | PFD1 | SC -met -lys |
| Scer | Scer-intralogous | GIM5 F[1,2] | YKE2 F[3] | GIM5 | YKE2 | SC -met -lys |
| Scer | Scer-intralogous | PAC10 F[1,2] | GIM3 F[3] | PAC10 | GIM3 | SC -met -lys |
| Scer | Scer-intralogous | PAC10 F[1,2] | GIM4 F[3] | PAC10 | GIM4 | SC -met -lys |
| Scer | Scer-intralogous | PAC10 F[1,2] | GIM5 F[3] | PAC10 | GIM5 | SC -met -lys |
| Scer | Scer-intralogous | PAC10 F[1,2] | CAB029 | PAC10 | PFD1 | SC -met -lys |
| Scer | Scer-intralogous | PAC10 F[1,2] | YKE2 F[3] | PAC10 | YKE2 | SC -met -lys |
| Scer | Scer-intralogous | PFD1 F[1,2] | GIM3 F[3] | PFD1 | GIM3 | SC -met -lys |
| Scer | Scer-intralogous | PFD1 F[1,2] | GIM4 F[3] | PFD1 | GIM4 | SC -met -lys |
| Scer | Scer-intralogous | PFD1 F[1,2] | GIM5 F[3] | PFD1 | GIM5 | SC -met -lys |
| Scer | Scer-intralogous | PFD1 F[1,2] | PAC10 F[3] | PFD1 | PAC10 | SC -met -lys |
| Scer | Scer-intralogous | PFD1 F[1,2] | YKE2 F[3] | PFD1 | YKE2 | SC -met -lys |
| Scer | Scer-intralogous | YKE2 F[1,2] | GIM3 F[3] | YKE2 | GIM3 | SC -met -lys |
| Scer | Scer-intralogous | YKE2 F[1,2] | GIM4 F[3] | YKE2 | GIM4 | SC -met -lys |
| Scer | Scer-intralogous | YKE2 F[1,2] | GIM5 F[3] | YKE2 | GIM5 | SC -met -lys |
| Scer | Scer-intralogous | YKE2 F[1,2] | PAC10 F[3] | YKE2 | PAC10 | SC -met -lys |
| Scer | Scer-intralogous | YKE2 F[1,2] | CAB029 | YKE2 | PFD1 | SC -met -lys |
| Scer | Scer-intralogous | CAB052 | BY4742 | GIM3 | GIM4 | SC -met -lys |
| Scer | Scer-intralogous | CAB053 | BY4742 | GIM3 | GIM5 | SC -met -lys |
| Scer | Scer-intralogous | CAB054 | BY4742 | GIM3 | PAC10 | SC -met -lys |
| Scer | Scer-intralogous | CAB055 | BY4742 | GIM3 | YKE2 | SC -met -lys |
| Scer | Scer-intralogous | CAB056 | BY4742 | GIM4 | GIM5 | SC -met -lys |
| Scer | Scer-intralogous | CAB057 | BY4742 | GIM4 | PAC10 | SC -met -lys |
| Scer | Scer-intralogous | CAB058 | BY4742 | GIM4 | YKE2 | SC -met -lys |
| Scer | Scer-intralogous | CAB059 | BY4742 | GIM5 | PAC10 | SC -met -lys |
| Scer | Scer-intralogous | CAB060 | BY4742 | GIM5 | YKE2 | SC -met -lys |
| Scer | Scer-intralogous | CAB061 | BY4742 | PAC10 | YKE2 | SC -met -lys |
| Scer | Scer-intralogous | CAB047 | BY4742 | PFD1 | GIM3 | SC -met -lys |
| Scer | Scer-intralogous | CAB048 | BY4742 | PFD1 | GIM4 | SC -met -lys |
| Scer | Scer-intralogous | CAB049 | BY4742 | PFD1 | GIM5 | SC -met -lys |
| Scer | Scer-intralogous | CAB050 | BY4742 | PFD1 | PAC10 | SC -met -lys |
| Scer | Scer-intralogous | CAB051 | BY4742 | PFD1 | YKE2 | SC -met -lys |
| Hybrid | Scer-intralogous | CAB052 | MG031 | GIM3 | GIM4 | SC -met -ade |
| Hybrid | Scer-intralogous | CAB053 | MG031 | GIM3 | GIM5 | SC -met -ade |
| Hybrid | Scer-intralogous | CAB054 | MG031 | GIM3 | PAC10 | SC -met -ade |
| Hybrid | Scer-intralogous | CAB055 | MG031 | GIM3 | YKE2 | SC -met -ade |
| Hybrid | Scer-intralogous | CAB056 | MG031 | GIM4 | GIM5 | SC -met -ade |
| Hybrid | Scer-intralogous | CAB057 | MG031 | GIM4 | PAC10 | SC -met -ade |
| Hybrid | Scer-intralogous | CAB058 | MG031 | GIM4 | YKE2 | SC -met -ade |
| Hybrid | Scer-intralogous | CAB059 | MG031 | GIM5 | PAC10 | SC -met -ade |
| Hybrid | Scer-intralogous | CAB060 | MG031 | GIM5 | YKE2 | SC -met -ade |
| Hybrid | Scer-intralogous | CAB061 | MG031 | PAC10 | YKE2 | SC -met -ade |

| | | | | | | |
|---|---|---|---|---|---|---|
| Hybrid | Scer-intralogous | CAB047 | MG031 | PFD1 | GIM3 | SC -met -ade |
| Hybrid | Scer-intralogous | CAB048 | MG031 | PFD1 | GIM4 | SC -met -ade |
| Hybrid | Scer-intralogous | CAB049 | MG031 | PFD1 | GIM5 | SC -met -ade |
| Hybrid | Scer-intralogous | CAB050 | MG031 | PFD1 | PAC10 | SC -met -ade |
| Hybrid | Scer-intralogous | CAB051 | MG031 | PFD1 | YKE2 | SC -met -ade |
| Hybrid | Interlogous | GIM3 F[1,2] | CAB040 | GIM3 | GIM4 | SC -met -ade |
| Hybrid | Interlogous | GIM3 F[1,2] | CAB041 | GIM3 | GIM5 | SC -met -ade |
| Hybrid | Interlogous | GIM3 F[1,2] | CAB042 | GIM3 | PAC10 | SC -met -ade |
| Hybrid | Interlogous | GIM3 F[1,2] | CAB038 | GIM3 | PFD1 | SC -met -ade |
| Hybrid | Interlogous | GIM3 F[1,2] | CAB043 | GIM3 | YKE2 | SC -met -ade |
| Hybrid | Interlogous | GIM4 F[1,2] | CAB039 | GIM4 | GIM3 | SC -met -ade |
| Hybrid | Interlogous | GIM4 F[1,2] | CAB041 | GIM4 | GIM5 | SC -met -ade |
| Hybrid | Interlogous | GIM4 F[1,2] | CAB042 | GIM4 | PAC10 | SC -met -ade |
| Hybrid | Interlogous | GIM4 F[1,2] | CAB038 | GIM4 | PFD1 | SC -met -ade |
| Hybrid | Interlogous | GIM4 F[1,2] | CAB043 | GIM4 | YKE2 | SC -met -ade |
| Hybrid | Interlogous | GIM5 F[1,2] | CAB039 | GIM5 | GIM3 | SC -met -ade |
| Hybrid | Interlogous | GIM5 F[1,2] | CAB040 | GIM5 | GIM4 | SC -met -ade |
| Hybrid | Interlogous | GIM5 F[1,2] | CAB042 | GIM5 | PAC10 | SC -met -ade |
| Hybrid | Interlogous | GIM5 F[1,2] | CAB038 | GIM5 | PFD1 | SC -met -ade |
| Hybrid | Interlogous | GIM5 F[1,2] | CAB043 | GIM5 | YKE2 | SC -met -ade |
| Hybrid | Interlogous | PAC10 F[1,2] | CAB039 | PAC10 | GIM3 | SC -met -ade |
| Hybrid | Interlogous | PAC10 F[1,2] | CAB040 | PAC10 | GIM4 | SC -met -ade |
| Hybrid | Interlogous | PAC10 F[1,2] | CAB041 | PAC10 | GIM5 | SC -met -ade |
| Hybrid | Interlogous | PAC10 F[1,2] | CAB038 | PAC10 | PFD1 | SC -met -ade |
| Hybrid | Interlogous | PAC10 F[1,2] | CAB043 | PAC10 | YKE2 | SC -met -ade |
| Hybrid | Interlogous | PFD1 F[1,2] | CAB039 | PFD1 | GIM3 | SC -met -ade |
| Hybrid | Interlogous | PFD1 F[1,2] | CAB040 | PFD1 | GIM4 | SC -met -ade |
| Hybrid | Interlogous | PFD1 F[1,2] | CAB041 | PFD1 | GIM5 | SC -met -ade |
| Hybrid | Interlogous | PFD1 F[1,2] | CAB042 | PFD1 | PAC10 | SC -met -ade |
| Hybrid | Interlogous | PFD1 F[1,2] | CAB043 | PFD1 | YKE2 | SC -met -ade |
| Hybrid | Interlogous | YKE2 F[1,2] | CAB039 | YKE2 | GIM3 | SC -met -ade |
| Hybrid | Interlogous | YKE2 F[1,2] | CAB040 | YKE2 | GIM4 | SC -met -ade |
| Hybrid | Interlogous | YKE2 F[1,2] | CAB041 | YKE2 | GIM5 | SC -met -ade |
| Hybrid | Interlogous | YKE2 F[1,2] | CAB042 | YKE2 | PAC10 | SC -met -ade |
| Hybrid | Interlogous | YKE2 F[1,2] | CAB038 | YKE2 | PFD1 | SC -met -ade |
| Hybrid | Interlogous | CAB020 | GIM4 F[3] | GIM3 | GIM4 | SC -trp -leu |
| Hybrid | Interlogous | CAB020 | GIM5 F[3] | GIM3 | GIM5 | SC -trp -leu |
| Hybrid | Interlogous | CAB020 | PAC10 F[3] | GIM3 | PAC10 | SC -trp -leu |
| Hybrid | Interlogous | CAB020 | CAB029 | GIM3 | PFD1 | SC -trp -leu |
| Hybrid | Interlogous | CAB020 | YKE2 F[3] | GIM3 | YKE2 | SC -trp -leu |
| Hybrid | Interlogous | CAB021 | GIM3 F[3] | GIM4 | GIM3 | SC -trp -leu |
| Hybrid | Interlogous | CAB021 | GIM5 F[3] | GIM4 | GIM5 | SC -trp -leu |
| Hybrid | Interlogous | CAB021 | PAC10 F[3] | GIM4 | PAC10 | SC -trp -leu |
| Hybrid | Interlogous | CAB021 | CAB029 | GIM4 | PFD1 | SC -trp -leu |
| Hybrid | Interlogous | CAB021 | YKE2 F[3] | GIM4 | YKE2 | SC -trp -leu |

| | | | | | | |
|---|---|---|---|---|---|---|
| Hybrid | Interlogous | CAB022 | GIM3 F[3] | GIM5 | GIM3 | SC -trp -leu |
| Hybrid | Interlogous | CAB022 | GIM4 F[3] | GIM5 | GIM4 | SC -trp -leu |
| Hybrid | Interlogous | CAB022 | PAC10 F[3] | GIM5 | PAC10 | SC -trp -leu |
| Hybrid | Interlogous | CAB022 | CAB029 | GIM5 | PFD1 | SC -trp -leu |
| Hybrid | Interlogous | CAB022 | YKE2 F[3] | GIM5 | YKE2 | SC -trp -leu |
| Hybrid | Interlogous | CAB023 | GIM3 F[3] | PAC10 | GIM3 | SC -trp -leu |
| Hybrid | Interlogous | CAB023 | GIM4 F[3] | PAC10 | GIM4 | SC -trp -leu |
| Hybrid | Interlogous | CAB023 | GIM5 F[3] | PAC10 | GIM5 | SC -trp -leu |
| Hybrid | Interlogous | CAB023 | CAB029 | PAC10 | PFD1 | SC -trp -leu |
| Hybrid | Interlogous | CAB023 | YKE2 F[3] | PAC10 | YKE2 | SC -trp -leu |
| Hybrid | Interlogous | CAB046 | GIM3 F[3] | PFD1 | GIM3 | SC -trp -leu |
| Hybrid | Interlogous | CAB046 | GIM4 F[3] | PFD1 | GIM4 | SC -trp -leu |
| Hybrid | Interlogous | CAB046 | GIM5 F[3] | PFD1 | GIM5 | SC -trp -leu |
| Hybrid | Interlogous | CAB046 | PAC10 F[3] | PFD1 | PAC10 | SC -trp -leu |
| Hybrid | Interlogous | CAB046 | YKE2 F[3] | PFD1 | YKE2 | SC -trp -leu |
| Hybrid | Interlogous | CAB024 | GIM3 F[3] | YKE2 | GIM3 | SC -trp -leu |
| Hybrid | Interlogous | CAB024 | GIM4 F[3] | YKE2 | GIM4 | SC -trp -leu |
| Hybrid | Interlogous | CAB024 | GIM5 F[3] | YKE2 | GIM5 | SC -trp -leu |
| Hybrid | Interlogous | CAB024 | PAC10 F[3] | YKE2 | PAC10 | SC -trp -leu |
| Hybrid | Interlogous | CAB024 | CAB029 | YKE2 | PFD1 | SC -trp -leu |
| Hybrid | Intralogous Suva | CAB067 | BY4742 | GIM3 | GIM4 | SC -trp -leu |
| Hybrid | Intralogous Suva | CAB068 | BY4742 | GIM3 | GIM5 | SC -trp -leu |
| Hybrid | Intralogous Suva | CAB069 | BY4742 | GIM3 | PAC10 | SC -trp -leu |
| Hybrid | Intralogous Suva | CAB070 | BY4742 | GIM3 | YKE2 | SC -trp -leu |
| Hybrid | Intralogous Suva | CAB071 | BY4742 | GIM4 | GIM5 | SC -trp -leu |
| Hybrid | Intralogous Suva | CAB072 | BY4742 | GIM4 | PAC10 | SC -trp -leu |
| Hybrid | Intralogous Suva | CAB073 | BY4742 | GIM4 | YKE2 | SC -trp -leu |
| Hybrid | Intralogous Suva | CAB074 | BY4742 | GIM5 | PAC10 | SC -trp -leu |
| Hybrid | Intralogous Suva | CAB075 | BY4742 | GIM5 | YKE2 | SC -trp -leu |
| Hybrid | Intralogous Suva | CAB076 | BY4742 | PAC10 | YKE2 | SC -trp -leu |
| Hybrid | Intralogous Suva | CAB062 | BY4742 | PFD1 | GIM3 | SC -trp -leu |
| Hybrid | Intralogous Suva | CAB063 | BY4742 | PFD1 | GIM4 | SC -trp -leu |
| Hybrid | Intralogous Suva | CAB064 | BY4742 | PFD1 | GIM5 | SC -trp -leu |
| Hybrid | Intralogous Suva | CAB065 | BY4742 | PFD1 | PAC10 | SC -trp -leu |
| Hybrid | Intralogous Suva | CAB066 | BY4742 | PFD1 | YKE2 | SC -trp -leu |
| Suva | Intralogous Suva | CAB067 | MG031 | GIM3 | GIM4 | SC -trp -ade |
| Suva | Intralogous Suva | CAB068 | MG031 | GIM3 | GIM5 | SC -trp -ade |
| Suva | Intralogous Suva | CAB069 | MG031 | GIM3 | PAC10 | SC -trp -ade |
| Suva | Intralogous Suva | CAB070 | MG031 | GIM3 | YKE2 | SC -trp -ade |
| Suva | Intralogous Suva | CAB071 | MG031 | GIM4 | GIM5 | SC -trp -ade |
| Suva | Intralogous Suva | CAB072 | MG031 | GIM4 | PAC10 | SC -trp -ade |
| Suva | Intralogous Suva | CAB073 | MG031 | GIM4 | YKE2 | SC -trp -ade |
| Suva | Intralogous Suva | CAB074 | MG031 | GIM5 | PAC10 | SC -trp -ade |
| Suva | Intralogous Suva | CAB075 | MG031 | GIM5 | YKE2 | SC -trp -ade |
| Suva | Intralogous Suva | CAB076 | MG031 | PAC10 | YKE2 | SC -trp -ade |

| | | | | | | |
|---|---|---|---|---|---|---|
| Suva | Intralogous Suva | CAB062 | MG031 | PFD1 | GIM3 | SC -trp -ade |
| Suva | Intralogous Suva | CAB063 | MG031 | PFD1 | GIM4 | SC -trp -ade |
| Suva | Intralogous Suva | CAB064 | MG031 | PFD1 | GIM5 | SC -trp -ade |
| Suva | Intralogous Suva | CAB065 | MG031 | PFD1 | PAC10 | SC -trp -ade |
| Suva | Intralogous Suva | CAB066 | MG031 | PFD1 | YKE2 | SC -trp -ade |
| Suva | Intralogous Suva | CAB020 | CAB040 | GIM3 | GIM4 | SC -trp -ade |
| Suva | Intralogous Suva | CAB020 | CAB041 | GIM3 | GIM5 | SC -trp -ade |
| Suva | Intralogous Suva | CAB020 | CAB042 | GIM3 | PAC10 | SC -trp -ade |
| Suva | Intralogous Suva | CAB020 | CAB038 | GIM3 | PFD1 | SC -trp -ade |
| Suva | Intralogous Suva | CAB020 | CAB043 | GIM3 | YKE2 | SC -trp -ade |
| Suva | Intralogous Suva | CAB021 | CAB039 | GIM4 | GIM3 | SC -trp -ade |
| Suva | Intralogous Suva | CAB021 | CAB041 | GIM4 | GIM5 | SC -trp -ade |
| Suva | Intralogous Suva | CAB021 | CAB042 | GIM4 | PAC10 | SC -trp -ade |
| Suva | Intralogous Suva | CAB021 | CAB038 | GIM4 | PFD1 | SC -trp -ade |
| Suva | Intralogous Suva | CAB021 | CAB043 | GIM4 | YKE2 | SC -trp -ade |
| Suva | Intralogous Suva | CAB022 | CAB039 | GIM5 | GIM3 | SC -trp -ade |
| Suva | Intralogous Suva | CAB022 | CAB040 | GIM5 | GIM4 | SC -trp -ade |
| Suva | Intralogous Suva | CAB022 | CAB042 | GIM5 | PAC10 | SC -trp -ade |
| Suva | Intralogous Suva | CAB022 | CAB038 | GIM5 | PFD1 | SC -trp -ade |
| Suva | Intralogous Suva | CAB022 | CAB043 | GIM5 | YKE2 | SC -trp -ade |
| Suva | Intralogous Suva | CAB023 | CAB039 | PAC10 | GIM3 | SC -trp -ade |
| Suva | Intralogous Suva | CAB023 | CAB040 | PAC10 | GIM4 | SC -trp -ade |
| Suva | Intralogous Suva | CAB023 | CAB041 | PAC10 | GIM5 | SC -trp -ade |
| Suva | Intralogous Suva | CAB023 | CAB038 | PAC10 | PFD1 | SC -trp -ade |
| Suva | Intralogous Suva | CAB023 | CAB043 | PAC10 | YKE2 | SC -trp -ade |
| Suva | Intralogous Suva | CAB046 | CAB039 | PFD1 | GIM3 | SC -trp -ade |
| Suva | Intralogous Suva | CAB046 | CAB040 | PFD1 | GIM4 | SC -trp -ade |
| Suva | Intralogous Suva | CAB046 | CAB041 | PFD1 | GIM5 | SC -trp -ade |
| Suva | Intralogous Suva | CAB046 | CAB042 | PFD1 | PAC10 | SC -trp -ade |
| Suva | Intralogous Suva | CAB046 | CAB043 | PFD1 | YKE2 | SC -trp -ade |
| Suva | Intralogous Suva | CAB024 | CAB039 | YKE2 | GIM3 | SC -trp -ade |
| Suva | Intralogous Suva | CAB024 | CAB040 | YKE2 | GIM4 | SC -trp -ade |
| Suva | Intralogous Suva | CAB024 | CAB041 | YKE2 | GIM5 | SC -trp -ade |
| Suva | Intralogous Suva | CAB024 | CAB042 | YKE2 | PAC10 | SC -trp -ade |
| Suva | Intralogous Suva | CAB024 | CAB038 | YKE2 | PFD1 | SC -trp -ade |

### Table S2. Composition of the different media used in this study.

| Medium | Composition |
|---|---|
| YPD | 1 % yeast extract |
| | 2 % tryptone |
| | 2 % glucose |
| | 2 % agar* |
| NAT | YPD + |
| | 100 µg/ml of Nourseothricin (NAT) |
| HYG | YPD + |
| | 250 µg/ml of Hygromycin (HygB) |
| NAT+HygB | YPD + |
| | 100 µg/ml NAT |
| | 250 µg/ml HygB |
| SC (Synthetic Complete) | 0.174 % yeast nitrogen base without amino acid without ammonium |
| | 2 % glucose |
| | 0.1 % L-glutamic acid |
| | Complete amino acid dropout mix |
| | 2 % agar* |
| SC -lys | SC + |
| | Amino acid dropout mix lacking lysine (instead of complete) |
| SC -met | SC + |
| | Amino acid dropout mix lacking methionine (instead of complete) |
| SC -met -lys | SC + |
| | Amino acid dropout mix lacking methionine and lysine (instead of complete) |
| SC -trp - ade | SC + |
| | Amino acid dropout mix lacking tryptophan and adenine (instead of complete) |
| SC -met - ade | SC + |
| | Amino acid dropout mix lacking methionine and adenine (instead of complete) |
| SC -trp -leu | SC + |
| | Amino acid dropout mix lacking tryptophan and leucine (instead of complete) |
| Enriched sporulation medium | 1 % potassium acetate |
| | 0.1 % yeast extract |
| | 0.05 % glucose |
| | 0.01 % amino-acid supplement powder mixture for sporulation |
| | 2 % agar |
| Amino-acid supplement powder mixture for sporulation | 2 g histidine |
| | 10 g leucine |
| | 2 g lysine |
| | 2 g uracil |
| MTX | 0.67 % yeast nitrogen base without amino acids and without ammonium sulfate |
| | 2 % glucose |
| | Amino acid drop-out without adenine |

| | |
|---|---|
| | 2.5 % noble agar |
| | 200 µg/ml methotrexate (MTX) diluted in DMSO |
| DMSO | Same as MTX with MTX replaced by DMSO |

* Agar is used for solid medium only

### Table S3. PCR primers used in the study.

| Name | Description | Sequence |
|---|---|---|
| O1-42 | Verification MATa – F | ACTCCACTTCAAGTAAGAGTTTG |
| O1-43 | Verification MATα – F | GCACGGAATATGGGACTACTTCG |
| O1-44 | Verification MATa/α – R | AGTCACATCAAGATCGTTTATGG |
| OP26-G4 | Universal primer F in *POP2* gene (*Saccharomyces* species) | ATGTTTCCTCAAGGGATGC |
| OP26-G7 | Universal primer R in *POP2* gene (*Saccharomyces* species) | TTGCTCAATTGGCAGAAGGA |
| OP151-A1 | Primer F to tag Suva *PFD1* gene with DHFR F amplified from pAG25-linker-F[1,2]-ADHterm, pAG32-linker-F[3]-ADHterm or pAG32-DHFR[3]-HPHNT1 | TGAAAAGACAATAGACAATCTAAAGGCAATGATGAAGAATggcggtggcg gatcaggaggc |
| OP151-B1 | Primer R to tag Suva *PFD1* gene with DHFR F amplified from pAG25-linker-F[1,2]-ADHterm or pAG32-linker-F[3]-ADHterm | TTTTCCCGTCCAAAAAACACTGATAGCTGCTACTCTTATAttcgacactg gatggcggcgttag |
| OP161-G6 | Primer R to tag Suva *PFD1* gene with DHFR F[3] amplified from pAG32-DHFR[3]-HPHNT1 | TTTTCCCGTCCAAAAAACACTGATAGCTGCTACTCTTATAgcataggcca ctagtggatc |
| OP151-C1 | Primer F to tag Suva *GIM3* gene with DHFR F amplified from pAG25-linker-F[1,2]-ADHterm, pAG32-linker-F[3]-ADHterm or pAG32-linker-F[3]-ADHterm | CCTGTACGCCAAGTTTGGCGATAATATCAATCTTGAACGTggcggtggcg gatcaggaggc |
| OP151 D1 | Primer R to tag Suva *GIM3* gene with DHFR F amplified from pAG25-linker-F[1,2]-ADHterm or pAG32-linker-F[3]-ADHterm | TTTGTGGAGACACCCACAGCCCACACAGCAGGGTAGGGCCttcgacactg gatggcggcgttag |
| OP161-G7 | Primer R to tag Suva *GIM3* gene with DHFR F[3] amplified from pAG32-DHFR[3]-HPHNT1 | TTTGTGGAGACACCCACAGCCCACACAGCAGGGTAGGGCCgcataggcca ctagtggatc |
| OP151-E1 | Primer F to tag Suva *GIM4* gene with DHFR F amplified from pAG25-linker-F[1,2]-ADHterm, pAG32-linker-F[3]-ADHterm or pAG32-linker-F[3]-ADHterm | GAAATGGAAAAAAGACAACAAAATTCAAGTTGTCAAAAACggcggtggcg gatcaggaggc |
| OP151-F1 | Primer R to tag Suva *GIM4* gene with DHFR F amplified from pAG25-linker-F[1,2]-ADHterm or pAG32-linker-F[3]-ADHterm | CTGCTAGACAAAGTATATCAAAAAAGTTAGAATAATTAGTttcgacactg gatggcggcgttag |
| OP161-G8 | Primer R to tag Suva *GIM4* gene with DHFR F[3] amplified from pAG32-DHFR[3]-HPHNT1 | CTGCTAGACAAAGTATATCAAAAAAGTTAGAATAATTAGTgcataggcca ctagtggatc |
| OP151-G1 | Primer F to tag Suva *GIM5* gene with DHFR F amplified from pAG25-linker-F[1,2]-ADHterm, pAG32-linker-F[3]-ADHterm or pAG32-linker-F[3]-ADHterm | GCAGCAGCAACAGCAGCAGCAGAAAGAGGCTACCACAGCCggcggtggcg gatcaggaggc |
| OP151-H1 | Primer R to tag Suva *GIM5* gene with DHFR F amplified from pAG25-linker-F[1,2]-ADHterm or pAG32-linker-F[3]-ADHterm | ATAACGAAGAGATATAAAGGAAGTATAGATAACAGATGTAttcgacactg gatggcggcgttag |
| OP161-G9 | Primer R to tag Suva *GIM5* gene with DHFR F[3] amplified from pAG32-DHFR[3]-HPHNT1 | ATAACGAAGAGATATAAAGGAAGTATAGATAACAGATGTAgcataggcca ctagtggatc |
| OP151-A2 | Primer F to tag Suva *PAC10* gene with DHFR F amplified from pAG25-linker-F[1,2]-ADHterm, pAG32-linker-F[3]-ADHterm or pAG32-linker-F[3]-ADHterm | ATTGAAGCAGGCCGAGGAAGGGACCCAAAATCTGAAGATAggcggtggcg gatcaggaggc |
| OP151-B2 | Primer R to tag Suva *PAC10* gene with DHFR F amplified from pAG25-linker-F[1,2]-ADHterm or pAG32-linker-F[3]-ADHterm | TAGTACTGGCTTCTAGTGGGGTCGGGTGCCAAAACGGCTGttcgacactg gatggcggcgttag |
| OP161-G10 | Primer R to tag Suva *PAC10* gene with DHFR F[3] amplified from pAG32-DHFR[3]-HPHNT1 | TAGTACTGGCTTCTAGTGGGGTCGGGTGCCAAAACGGCTGgcataggcca ctagtggatc |
| OP151-C2 | Primer F to tag Suva *YKE2* gene with DHFR F amplified from pAG25-linker-F[1,2]-ADHterm, pAG32-linker-F[3]-ADHterm or pAG32-linker-F[3]-ADHterm | CAAACTAAATAATGCCGCAACTGCCGCTGGCCCAGACAGGggcggtggcg gatcaggaggc |
| OP151-D2 | Primer R to tag Suva *YKE2* gene with DHFR F amplified from pAG25-linker-F[1,2]-ADHterm or pAG32-linker-F[3]-ADHterm | GTATGTTTTCTGTTCTTCTTTCAACATAATGCAATTTCTTttcgacactg gatggcggcgttag |
| OP161-G11 | Primer R to tag Suva *YKE2* gene with DHFR F[3] amplified from pAG32-DHFR[3]-HPHNT1 | GTATGTTTTCTGTTCTTCTTTCAACATAATGCAATTTCTTgcataggcca ctagtggatc |
| OP151-C3 | Primer F to tag Scer *PFD1* gene with DHFR F amplified from pAG25-linker-F[1,2]-ADHterm or pAG32-linker-F[3]-ADHterm | TGAAAAAACAATAGACAATCTAAAGGCATTGATGAAGAATggcggtggcg gatcaggaggc |
| OP151-D3 | Primer R to tag Scer *PFD1* gene with DHFR F amplified from pAG25-linker-F[1,2]-ADHterm or pAG32-linker-F[3]-ADHterm | GTTTTCCGGCTAAGAAAGGAAAGGCTATTGCCGCTTTCTTttcgacactg gatggcggcgttag |
| OP151-E2 | Primer F at the 3' extremity to confirm tagging of Suva *PFD1* | GGAGATCCTGTGGAAAATCG |
| OP151-F2 | Primer F at the 3' extremity to confirm tagging of Suva *GIM3* | GCACAGCTCGAAGAAGATGT |
| OP151-G2 | Primer F at the 3' extremity to confirm tagging of Suva *GIM4* | GGGCCATGATAAAGATGAGC |
| OP151-H2 | Primer F at the 3' extremity to confirm tagging of Suva *GIM5* | GGAACCGGCTATTACGTTGA |
| OP151-A3 | Primer F at the 3' extremity to confirm tagging of Suva *PAC10* | CATCGAGCTGTTGCAGAAGA |
| OP151-B3 | Primer F at the 3' extremity to confirm tagging of Suva *YKE2* | CCAGTGGAACAAAGTGAAGC |

| | | |
|---|---|---|
| OP151-E3 | Primer F at the 3' extremity to confirm tagging of Scer PFD1 | TGACGAAACTGTTCTTCTGG |
| OP151-F3 | Primer F at the 3' extremity to confirm tagging of Scer GIM3 | AGAGACTTTAGAGGACAAGC |
| OP151-G3 | Primer F at the 3' extremity to confirm tagging of Scer GIM4 | TACAGGATGATAGGTGGCGC |
| OP151-H3 | Primer F at the 3' extremity to confirm tagging of Scer GIM5 | TTCCTTGTCCATCGAGGCCC |
| OP151-A4 | Primer F at the 3' extremity to confirm tagging of Scer PAC10 | GGACGTAGAGTTTTTGAGGG |
| OP151-B4 | Primer F at the 3' extremity to confirm tagging of Scer YKE2 | ATGCGAAAAGAACATAAGGG |
| O1-50 | Primer R in ADHterm to confirm DHFR tagging | CCATCTTTTCGTAAATTTCTG |

***Supplementary data legends***

**Data S1.** Elution profiles of proteins obtained from SEC-PCP-SILAC experiment.

**Data S2.** Interaction scores of the proteins obtained from SEC-PCP-SILAC experiment.

**Data S3.** Protein abundance and dosage balance scores obtained from SEC-PCP-SILAC experiment.

**Data S4.** Interaction scores of the proteins within the prefoldin complex obtained from DHFR-PCA experiment.

**Data S5.** Interaction scores ratios for the comparison of PPIs in hybrids with those in parents.

**Data S6.** Gene set enrichment analysis for the comparison of PPIs in hybrids with those in parents.
